# A single-stranded DNA virus replicates in the mitochondria of a marine oomycete

**DOI:** 10.64898/2026.04.22.720165

**Authors:** Kohei Sakuta, Mart Krupovic, Ondrej Hejna, Cristiana Maia, Marília Horta Jung, Ken Komatsu, Hiromitsu Moriyama, Thomas Jung, Leticia Botella

## Abstract

A virus with a circular single-stranded DNA genome, HthCRESSV1, was discovered in an oomycete, *Halophytophthora thermoambigua*, isolated from brackish waters off the Algarve coast in southern Portugal. Phylogenetic analyses place this virus outside of all currently classified families within the phylum *Cressdnaviricota*, suggesting it represents a distinct lineage. Cellular fractionation, mitochondrial marker co-enrichment, and Southern blot analyses indicate that HthCRESSV1 is associated with mitochondria and replicates episomally. The virus is stably transmitted through zoospores and, notably, infected host isolates are associated with reduced growth and changes in temperature-dependent performance. Homologous sequences corresponding to both the replication-associated protein and a membrane-associated hypothetical protein were identified in the mitochondrial genomes of several oomycete species, suggesting recurrent endogenization of HthCRESSV1-like viruses in oomycete mitochondria. Together, these findings expand the known diversity of oomycete-associated DNA viruses, identify mitochondria as an unexpected niche for DNA virus replication, and highlight marine environments as underexplored reservoirs of DNA viruses in stramenopiles.

## INTRODUCTION

Metagenomic analyses have revealed that viruses with single-stranded DNA (ssDNA) genomes are ubiquitous, remarkably diverse, and infect cellular organisms across the tree of life [**1–3**]. One of the most diverse groups of ssDNA viruses is represented by members of the phylum *Cressdnaviricota* [**4**], formerly known as the circular, Rep-encoding single-stranded DNA (CRESS DNA) viruses [**1**]. These viruses share the characteristic two-domain rolling circle replication (RCR) initiation protein (Rep) comprising the N-terminal HUH superfamily endonuclease domain and the C-terminal superfamily 3 helicase domain [**5**]. All cressdnaviricots characterized thus far build non-enveloped, icosahedral capsids and encode CPs with the jelly-roll fold (antiparallel 8-strand β-barrel) [**4, 6**]. Whereas the Rep is universally encoded across *Cressdnaviricota*, the capsid protein (CP) genes of these viruses display considerable sequence divergence and in some lineages have been demonstrated to undergo gene replacement with non-orthologous, but structurally related CPs from other viruses [**7–10**]. Thus, the phylogenetic analysis of the Rep proteins is used as a framework for the classification of cressdnaviricots into families, orders and classes.

Currently, *Cressdnaviricota* comprises two classes (*Arfiviricetes* and *Respensiviricetes*), 13 orders and 24 families [**11**]. All cressdnaviricots for which the hosts have been experimentally determined infect eukaryotes, including plants (*i.e., Nanoviridae*, *Geminiviridae*, *Amesuviridae*, *Metaxyviridae*), fungi (*Genomoviridae*), animals (*i.e., Circoviridae*, *Redondoviridae*), algae (*Bacilladnaviridae*) and protists (*i.e., Naryaviridae*, *Nenyaviridae*, *Vilyaviridae, Oomyviridae*) [**12–14**]. However, hosts of viruses representing a considerable fraction of families are currently unknown. Furthermore, many ssDNA viruses, especially those discovered through metagenomics, remain unclassified. Thus, the extent of cressdnaviricot diversity and their distribution in eukaryotes remain to be fully appreciated.

In unicellular eukaryotes, ssDNA viruses have been described for several economically important plant pathogenic filamentous fungi, including *Sclerotinia sclerotiorum* (infected by Sclerotinia sclerotiorum hypovirulence-associated DNA virus 1; SsHADV1), *Fusarium graminearum* (Fusarium graminearum gemytripvirus 1), *Diaporthe sojae* (Diaporthe sojae circular DNA virus 1), and *Botrytis cinerea* (Botrytis cinerea ssDNA virus 1) [**15–19**]. No ssDNA viruses infecting oomycetes have been isolated but metagenomic studies have uncovered the existence of multiple endogenous viral elements (EVEs), encompassing the Rep and CP genes typical of ssDNA viruses, in the genomes of oomycetes [**20**]. In particular, EVEs were found in chromosome 5 of *Phytophthora plurivora*, *Phytophthora parasitica* (synonym *Phytophthora nicotianae*), *Phytophthora fragariae*, *Aphanomyces astaci* and *Pythium insidiosum*. Furthermore, transcripts corresponding to ssDNA virus genes were detected in the transcriptomes of *Plasmopara halstedii* and *Aphanomyces invadans*, suggesting the presence of transcriptionally active viral elements and raising the possibility of ongoing or recent viral replication rather than solely degenerated genomic remnants [**20**]. Environmental metatranscriptomic analyses have further identified diverse cressdnaviricot sequences in various habitats known to harbor fungi and oomycetes, including soil, riverbed sediments, plant samples, and animal feces [**21–26**]. Therefore, viruses of fungi and oomycetes may play important, yet underappreciated roles in ecosystem structuring by controlling the behavior and dynamics of their hosts [**27**]. Indeed, several SsHADV1-like ssDNA viruses (family *Genomoviridae*) have been shown to induce hypovirulence of their phytopathogenic fungal hosts [**15, 16**].

In marine ecosystems, viral infections are observed in all organisms from bacteria to whales, and the presence of these viruses influences marine biological community composition and serves as a potential driving force in biogeochemical cycles [**28**]. Although dsDNA phages infecting bacteria are the most abundant viruses in marine ecosystems, metagenomic and metatranscriptomic studies have uncovered a substantial contribution to the global marine virome of ssDNA and RNA viruses that infect protists, including unicellular algae, fungi and fungus-like oomycetes [**29**]. RNA viruses have been identified from various host fungi infecting *Posidonia oceanica* (seagrass) and *Holothuria poli*, and oomycetes, including *Phytophthora condilina* and *Halophytophthora frigida* [**30–33**]. However, no DNA viruses infecting marine fungi or oomycetes have been identified to date.

The genus *Halophytophthora* constitutes a sister genus of the well-known plant pathogenic oomycete genus *Phytophthora*, comprising species with similar morphology and life cycles, almost exclusively inhabiting brackish and saltwater environments [**34**]. While some species, such as *Halophytophthora lateralis* (previously *Halophytophthora* sp. Zostera), have been demonstrated to be pathogenic to eelgrass (*Zostera marina*) [**35**], the genus *Halophytophthora* has been primarily considered as decomposers in mangrove ecosystems [**36, 37**]. The only report of viral infection in this genus is the co-infection of eight bunya-like viruses in *H. frigida* collected from estuaries in southern Portugal, and the effects of viral infection on the host remain unclear [**32**].

To expand the limited knowledge on viruses in *Halophytophthora*, we focused our viral discovery and characterization efforts on *Halophytophthora thermoambigua*, a species inhabiting marine and brackish-water environments [**38**]. *H. thermoambigua* emerged as a compelling candidate for virus screening due to two notable biological features: its self-sterile breeding system, with no observed oogonia, oospores, or antheridia, and its unusual phenotypic variability in colony morphology and optimal growth temperature among isolates (20, 25, or 27.5 °C). Here, we report the discovery of the first DNA virus identified in an oomycete. We demonstrate its mitochondrial localization, confirm its extrachromosomal replicative form, and provide evidence of stable vertical transmission. The virus is associated with reduced host growth and altered temperature-dependent performance.

## MATERIALS AND METHODS

### Strains and culture conditions

All *Halophytophthora* isolates/strains investigated in this study were collected in December 2015 from seven different marine and brackish water sites along the Algarve coast of southern Portugal using an in situ baiting method [**39**] in salt marshes, tidal pools, channels, lagoons, and estuaries that remained submerged even at low tide [**38**]. Supplementary Table S1 provides detailed information and species identification for all *Halophytophthora* isolates included in the experiments. All strains were cultured on salty V8-agar medium (sV8A; 16 g agar, 3 g CaCO3, 100 mL Campbell’s V8 juice, 450 mL distilled water, 450 mL seawater; [**38**] or liquid V8-medium containing seawater adjusted to 50% concentration (sV8).

### RNA extraction

Total RNA was purified from approximately 100 mg of 7-10-days old mycelium using RNAzol® RT Column Kit [**40**] and treated with TURBO DNA-free™ Kit (Ambion). RNA was quantified in Qubit® 2.0 Fluorometer (Invitrogen), and quality was tested by Tape Station 4200 (Agilent). One RNA pool was prepared with eight isolates of the ex-type species of *Halophytophthora* described in Maia et al. [**38**].

### RNA library preparation and total stranded RNA sequencing

Around 1 μg of total RNA, eluted in RNase-free water, was sent to IABio in Olomouc, Czech Republic, for RNA library construction and deep sequencing. RNA was depleted using the NEBNext rRNA Depletion Kit (Human/Mouse/Rat) and prepared with the NEBNext Ultra II Directional RNA Library Prep Kit for Illumina, along with NEBNext Multiplex Oligos for Illumina (Unique Dual Index Primer Pairs). Library quality control was evaluated with the Agilent Bioanalyzer 2100 High Sensitivity DNA Kit. The KAPA Library Quantification Kit for the Illumina platform facilitated absolute qPCR-based quantification of the Illumina libraries, flanked by the P5 and P7 flow cell oligo sequences. The libraries underwent paired-end (PE) (2 × 150 nt) sequencing on a NovaSeq6000 (DS-150) from Illumina, San Diego, CA, USA, using the NovaSeq S4 v1.5 reagent kit. An “in-lane” PhiX control spike was included in each lane of the flow cell.

### Virus identification bioinformatics pipeline

#### Preprocessing of RNA-seq data

Raw data were processed by BaseSpace cloud interface (Illumina) in default settings. The processing was carried out on the local server of the University of South Bohemia in České Budějovice, Czech Republic. The basecalling, adapter clipping, and quality filtering were carried out using Bcl2fastq v2.20.0.422 Conversion Software (Illumina). For the initial quality assessment, we used FastQC v0.11.9 (https://www.bioinformatics.babraham.ac.uk/projects/fastqc/). The adapter sequences, provided by the sequencing firm, were verified using BBTools v37.87 (https://jgi.doe.gov/data-and-tools/software-tools/bbtools/) [**41**]. Following processing with Cutadapt v4.7 (https://cutadapt.readthedocs.io/en/stable/) [**42**] involved trimming of N bases, adapter sequences, and low-quality ends (<30). Reads falling short of 50 bp were discarded from subsequent analyses. Post-trimming quality was again re-assessed with FastQC. Remaining rRNA reads were removed using SortMeRNA v4.3.6 (https://github.com/sortmerna/sortmerna/) [**43**]. SILVA SSU and LSU databases filtered to include only Oomycota rRNA sequences were used as references (https://www.arb-silva.de/).

#### Alignment/Read mapping to host genome

For mapping reads to the host genome, the STAR v2.7.9a [**44**] program was employed with default settings. Two genomes, *Halophytophthora batemanensis* (GCA_023338225.1) and *Halophytophthora polymorphica* (GCA_023338205.1) were used as references. After alignment to the reference sequence, all mapped reads were discarded and only unmapped reads were utilized further.

#### De novo assembly of unmapped reads for detection of novel viruses

The unmapped reads were used for de novo assembly. To assemble these reads into contigs, the SPAdes program v3.15.5 (https://github.com/ablab/spades/) [**45**] was used with default settings for metagenomics. For further analysis, assembled contigs shorter than 500 bp were discarded.

#### Search for similarity in virus databases

Final contigs were cross-referenced against several databasesusing the Basic Local Alignment Search Tool (BLAST) v2.10.0 (https://blast.ncbi.nlm.nih.gov/doc/blast-help/downloadblastdata.html/ [**46**]. As a reference, we used viral reference sequences from NCBI. BLASTn was applied for the nucleotide database accessible at https://www.ncbi.nlm.nih.gov/labs/virus/vssi/#/virus?SeqType_s=Nucleotide. Similarly, BLASTx was employed for the protein database available at https://www.ncbi.nlm.nih.gov/labs/virus/vssi/#/virus?SeqType_s=Protein, and also to search the UniProt database for a specified virus taxon at https://www.uniprot.org/uniprot/?query=taxonomy:10239. Contigs with an e-value higher than 1e-3 were discarded. Subsequently, the remaining contigs were processed in a similar manner using the entire range of databases. For BLASTn, the NCBI nucleotide (nt) database from NCBI, available at https://ftp.ncbi.nlm.nih.gov/blast/db/nt*, was used. For BLASTx, both the NCBI nr database, accessible from at https://ftp.ncbi.nlm.nih.gov/blast/db/nr*, and the UniProt Knowledgebase (UniProtKB) available at https://www.uniprot.org/uniprotkb/, were employed. The BLAST search was restricted to the best hit per contig. The final list of candidate viral contigs was compiled by scanning the BLAST results for the search string ’vir’.

To identify homologous replicases and hypothetical proteins which might not be annotated in Blastx, a BLASTn search against the NCBI nt database using the full-length HthCRESSV1 genomic sequence as a query. Hits with high nucleotide similarity (e-value < 1e−10) were retrieved and manually inspected. Top matches corresponded to mitochondrial genome assemblies from diverse oomycete species. To confirm the genomic context and sequence identity, the HthCRESSV1-Rep and HP sequences were mapped to these mitochondrial genome assemblies using Geneious Prime (v2023.2.2), employing medium sensitivity and default parameters.

#### Coverage depth

The number of unique sequencing reads that align to each reference de novo assembled viral contig was calculated with the following formula: (Total reads mapped to the final identified virus * average read length)/virus genome or contig length) [**47**]. Coverage plots were visualised with Integrated Genome Viewer (IGV) tool and Geneious Prime® 2025.0.2. (Figure S1).

#### DNA extraction and PCR reactions

Purified DNA used for Southern hybridization was extracted using the modified cetyltrimethylammonium bromide (CTAB)-based method by Fukuhara et al. (1993). Specifically, approximately 0.1 g of dried mycelia harvested after 7 days of culture in sV8 liquid medium was used as starting material, and the extraction was performed on a small scale. DNA extraction from mycelia after single lesion isolation derived from zoospores was performed as described by Maia et al. [**38**]. DNA was stored at -80°C for long-term preservation. PCR reactions for complete sequence determination and detection of HtCRESSV1 were performed using the obtained DNA as template with primers listed in Table S2 and GoTaq (Promega, Wisconsin, USA).

#### Phylogenetic analyses

Maximum likelihood (ML) phylogenetic trees were constructed using RAxML-HPC v.8 on XSEDE via the CIPRES Science Gateway [**48**]. The analysis employed a rapid bootstrapping algorithm with the recommended parameters, and the GAMMA model was used to avoid extensive optimization of the best-scoring ML tree at the end of the run. The Jones–Taylor–Thornton (JTT) model was applied as the amino acid (aa) substitution model for protein sequences.

The resulting trees were visualized using iTOL v6 [**49**] (https://itol.embl.de).

#### Stem loop prediction

DNA secondary structure prediction and free energy calculations were carried out using two web-based computational tools. The mFold web server (hosted by UNAFold) was used to predict and visualize DNA secondary structures and determine associated thermodynamic parameters, including minimum free energy, under defined temperature and ionic conditions (https://www.unafold.org/mfold/applications/Structure-display-and-free-energy-determination.php). Additionally, the ViennaRNA Web Services platform (http://rna.tbi.univie.ac.at/) was also employed for further structure prediction and analysis. Although primarily designed for RNA, the ViennaRNA tools were adapted for DNA by specifying the appropriate nucleic acid type and parameters where applicable.

#### Analysis of the ORF2 protein sequence and structure

Secondary structure and putative transmembrane domains were predicted using the PSIPRED and MEMSAT-SVM algorithms, respectively, available through the PSIPRED Workbench [**50**]. Coiled-coil regions were predicted using PCOILS available through the MPI Bioinformatics Toolkit [**51**]. Structural modeling of ORF2 protein sequence was carried out using AlphaFold3 [**52**]. The models were built for different oligomeric states from monomer to homo-decamer. The quality of the models was evaluated by comparing their predicted template modelling (pTM) and the interface predicted template modelling (ipTM) scores. The resulting models were visualized using ChimeraX v1.11.1 [**53**].

#### Virus-like particle (VLP) purification and transmission electron microscopy (TEM) observation

Approximately 40 g of BD651 mycelium cultured in liquid s. V8 medium was collected using Miracloth, washed with distilled water, and incubated in Buffer A (0.1 M phosphate buffer, 0.2 M KCl) for 10 min. The mycelium was collected, homogenized in liquid nitrogen, and disrupted using a French press. Triton X-100 (1% final concentration) was added to the cell lysate, followed by stirring at 4°C for 30 min. The lysate was centrifuged at 1,000 *g* for 10 min to remove cellular debris. The supernatant was collected, mixed with an equal volume of chloroform, gently agitated for 30 min, and centrifuged at 8,000 *g* for 15 min. The aqueous phase was collected and centrifuged at 20,000 *g* for 30 min to pellet heavy membrane fractions. The resulting supernatant was ultracentrifuged at 100,000 *g* for 1 h using a Hitachi CP80WX P45AT rotor to pellet VLPs (P100,000). The pellet was resuspended in Buffer A and subjected to continuous sucrose density gradient centrifugation (10–40% w/v) at 80,000 *g* for 3 h at 4°C using a Hitachi CP80WX P28S swing rotor. Following ultracentrifugation, the gradient was fractionated from the top in 2-ml aliquots using a Hitachi DGF-U gradient fractionator, yielding 15 fractions. Total nucleic acids were extracted from each fraction, and HthCRESSV1-enriched fractions (fractions 4 and 5) were identified by HthCRESSV1-specific PCR. These fractions were pooled and concentrated by ultracentrifugation at 100,000 *g* for 1 h using a Hitachi CP80WX P45AT rotor. The pellet was resuspended in phosphate buffer, negatively stained with 2% uranyl acetate, and visualized by transmission electron microscopy (TEM) using a JEM-1400 Plus microscope (JEOL, Tokyo, Japan).

#### Southern Blot analysis

Approximately 5 µg of purified DNA extracted from strain BD651 were subjected to electrophoresis on a 0.8% agarose gel, followed by depurination treatment by immersing the gel in 0.25 M HCl and shaking for 5 min. After washing the gel with distilled water, DNA was denatured by shaking in denaturing solution (0.5 M NaOH, 1.5 M NaCl) for 15 min. This operation was performed twice. The denatured gel was then neutralized by shaking in neutralizing solution (0.5 M Tris-HCl, pH 7.5; 1.5 M NaCl) for 15 min, repeated twice, washed with distilled water, and equilibrated by incubation in 20× SSC for 10 min. Subsequently, the DNA was transferred to a positively charged nylon membrane (Roche, Basel, Switzerland) by capillary blotting. To detect the HtCRESSV1 genome, a DIG-labeled DNA probe was prepared using the HtCRESSV1-HP probe (nt 114-837) according to the protocol provided with the PCR DIG Labeling Mix (Roche, Basel, Switzerland). Hybridization with the DNA probe was performed at 50°C for 16 hours. Subsequent operations until detection were performed according to the method of Sakuta et al. [**54**].

#### Virus-like particle (VLP) purification and transmission electron microscopy (TEM) observation

Approximately 40 g of BD651 mycelium cultured in liquid sV8 medium was collected using Miracloth, washed with distilled water, and incubated in Buffer A (0.1 M phosphate buffer, 0.2 M KCl) for 10 min. The mycelium was collected, homogenized in liquid nitrogen, and disrupted using a French press. Triton X-100 (1% final concentration) was added to the cell lysate, followed by stirring at 4°C for 30 min. The lysate was centrifuged at 1,000 *g* for 10 min to remove cellular debris. The supernatant was collected, mixed with an equal volume of chloroform, gently agitated for 30 min, and centrifuged at 8,000 *g* for 15 min. The aqueous phase was collected and centrifuged at 20,000 *g* for 30 min to pellet heavy membrane fractions. The resulting supernatant was ultracentrifuged at 100,000 *g* for 1 h using a Hitachi CP80WX P45AT rotor to pellet VLPs (P100,000). The pellet was resuspended in Buffer A and subjected to continuous sucrose density gradient centrifugation (10–40% w/v) at 80,000 *g* for 3 h at 4°C using a Hitachi CP80WX P28S swing rotor. Following ultracentrifugation, the gradient was fractionated from the top in 2-ml aliquots using a Hitachi DGF-U gradient fractionator, yielding 15 fractions. Total nucleic acids were extracted from each fraction, and HthCRESSV1-enriched fractions (fractions 4 and 5) were identified by HthCRESSV1-specific PCR. These fractions were pooled and concentrated by ultracentrifugation at 100,000 *g* for 1 h using a Hitachi CP80WX P45AT rotor. The pellet was resuspended in phosphate buffer, negatively stained with 2% uranyl acetate, and visualized by TEM using a JEM-1400 Plus microscope (JEOL, Tokyo, Japan).

#### Mitochondrial purification and nucleic acid extraction

Approximately 20 g of mycelia were homogenized in 200 mL of isotonic solution (0.25 M sucrose, 100 mM Tris-HCl, pH 7.4, 10 mM EGTA). The homogenate was transferred to a 50 mL centrifuge tube and centrifuged at 1,000× g for 10 min to precipitate cell debris and nuclei (P1000). The supernatant was transferred to a Hizacks tube and centrifuged at 12,000× g for 15 min to obtain a crude mitochondrial fraction enriched in mitochondria (P12000). This supernatant was used as a membrane fraction and centrifuged at 100,000× g for 90 min using a P45AT rotor in a Hitachi CP80WX (Hitachi Koki, Japan), with the resulting pellet designated as the membrane fraction (P100000) and the supernatant as the cytosolic fraction (S100000). P12000 was purified using two rounds of sucrose density gradient centrifugation to obtain a more intact mitochondrial fraction. First, it was layered on a discontinuous sucrose solution prepared at concentrations of 15%–23%–32%–60% and centrifuged at 82,200× g for 3 hours using a P28S swing rotor in a Hitachi CP80WX (Hitachi Koki, Japan). After centrifugation, the purified mitochondrial concentrated fraction observed at the 32%–60% interface was carefully collected, diluted with isotonic solution, centrifuged at 10,000× g for 20 min, and the resulting pellet was resuspended in isotonic solution to obtain a semi-purified mitochondrial fraction. The second purification was performed by layering the semi-purified mitochondrial fraction on a sucrose solution prepared at 32%–60% concentration and ultracentrifuging under the same conditions as the first round. After centrifugation, 2 mL fractions were carefully collected from the top to obtain 15 fractions (F1–F15). Total DNA was extracted from the 15 fractions containing purified mitochondrial fractions using a modified method of Okada et al. [54]. Specifically, samples were mixed with 0.5 mL of 2× STE buffer (200 mM NaCl, 20 mM Tris-HCl pH 8.0, 2 mM EDTA pH 8.0) containing 1% SDS and 0.5 mL of 1:1 phenol/chloroform. The mixture was vortexed and centrifuged at 15,000 g for 5 min. The resulting supernatant was used as a total nucleic acid sample, and after ethanol precipitation, it was suspended in TE buffer containing RNase A and incubated at 37°C for 30 min to obtain a total DNA sample.

#### COX activity measurement

To measure the activity of cytochrome c oxidase, a mitochondrial marker enzyme, the Cytochrome c Oxidase Assay Kit (Sigma) was used according to the manufacturer’s instructions. Enzyme activity was measured by monitoring the absorbance at 550 nm for 1 minute, with the decrease in absorbance defined as ΔA and substituted into the following equation to measure enzyme activity: Units/mL·g = ΔA/min × 1.1 / (0.1 × 21.84 × X) (X: weight of source mycelia in g).

#### Zoospore induction and single lesion isolation, mycelial comparison and growth tests

Zoospore induction of *H. thermoambigua* was performed using modified protocols from Jesus et al. [**56, 57**]. Single lesion formation was achieved by floating young cork oak (*Quercus suber*) leaves on the water surface in non-sterile 50% seawater after zoospore induction and incubating at 20°C under natural light for 24 hours. The single lesions were excised with a razor blade and placed on PARPNH agar medium (V8 juice agar (V8A) supplemented with 10 µg/mL pimaricin, 200 µg/mL ampicillin, 10 µg/mL rifampicin, 25 µg/mL pentachloronitrobenzene (PCNB), 50 µg/mL nystatin, and 50 µg/mL hymexazol) to obtain single zoospore-derived isolates. The relationship between temperature and growth was examined by pre-culturing target strains on 90 mm V8A medium for one week to standardize the growth stage. Subsequently, three plates per isolate were incubated at 4, 8, 15, 20, 25, 28, 30, and 33°C in the dark. The test was conducted for 5 days or until the colonies nearly reached the edge of the Petri dish, and the average growth rate (cm/day) was calculated.

#### Attempted HthCRESSV1 elimination

A total of 20 single-zoospore strains derived from the isolate BD651 were prepared following the procedure described by Sakuta et al. [**58**] and Uchida et al. [**59**]. The regeneration medium was supplemented with ribavirin (300 µg ml^-1^ final concentration) and cycloheximide (5µg ml^-1^ final concentration), and 20 isolates were subsequently recovered from the regenerated mycelia. Total nucleic acids were then extracted from each culture grown on sV8 medium using the phenol-chloroform-isoamyl alcohol (PCI) method. The presence of the virus was finally verified by PCR employing a primer set specific to HtCRESSV1 (Table S2).

### Statistical Analysis

Colony diameter measurement data were subjected to analysis of variance. Dunnett’s test following one-way analysis of variance was applied for comparison of means between virus-infected and non-infected strain groups. Statistical significance was set at p < 0.05 (*p < 0.05, **p < 0.01).

## RESULTS

### Identification of Halophytophthora thermoambigua CRESS virus 1

Transcriptomic screening of eight pooled strains of *Halophytophthora* spp. (Table S1) yielded 729 sequencing reads that could be assembled into a 979 nt-long contig (NODE_2192_length_979; Figure S1) which displayed similarity to unclassified ssDNA viruses, including Tick-associated circular DNA virus (UTM74968.1; 27.6% identity, 76.3% query coverage, E=2.69E-11) and Grus japonensis CRESS-DNA-virus sp. (QTE03368.1; 26.9% identity, 71.2% coverage, E=1.17E-09). The hits to viruses encompassed a region encoding the Gemini_AL1 domain (Pfam: PF00799; E-value: 2.26e-07) corresponding to the catalytic domain of the geminiviral replication initiation protein (Rep), suggesting that the assembled contig corresponds to a previously unknown member of the phylum *Cressdnaviricota*. Notably, a BLASTx search queried with the contig sequence identified as the top hit a hypothetical protein (N0F65_000360) encoded by the mitochondrial genome of the oomycete *Lagenidium giganteum* (accession DAZ93511.1), with 67.81% identity and 45% query coverage (E-value=4e-63). The second and third best hits were also to two mitochondrially encoded hypothetical proteins from *Phytophthora sojae* (YP_001165403 and YP_001165402), pointing to the association of the identified virus with different oomycete species.

To identify the *Halophytophthora* strains infected with the putative ssDNA virus, total nucleic acids were extracted from the eight strains originally subjected to transcriptomic analysis, and virus identification was performed by PCR amplification using a pair of specific primers designed based on the contig sequence (Table S2). The positive signal was obtained exclusively with the DNA extracted from *H. thermoambigua* BD651, the ex-type strain of the species isolated from brackish water in Parque Natural da Ria Formosa, Santa Luzia, Tavira (Portugal). To obtain the complete genome of the putative ssDNA virus, DNA extracted from isolate BD651 was amplified by PCR using abutting primers (Table S2). After DNA purification, cloning, and sequencing, a complete circular genome of 2,793 nt was obtained (Figures 1A and S1) and denoted Halophytophthora thermoambigua circular rep-encoding single-stranded DNA virus 1 (HtCRESSV1) (GenBank accession number PV883009, SRA Bioproject PRJNA1437999).

**Figure 1.**
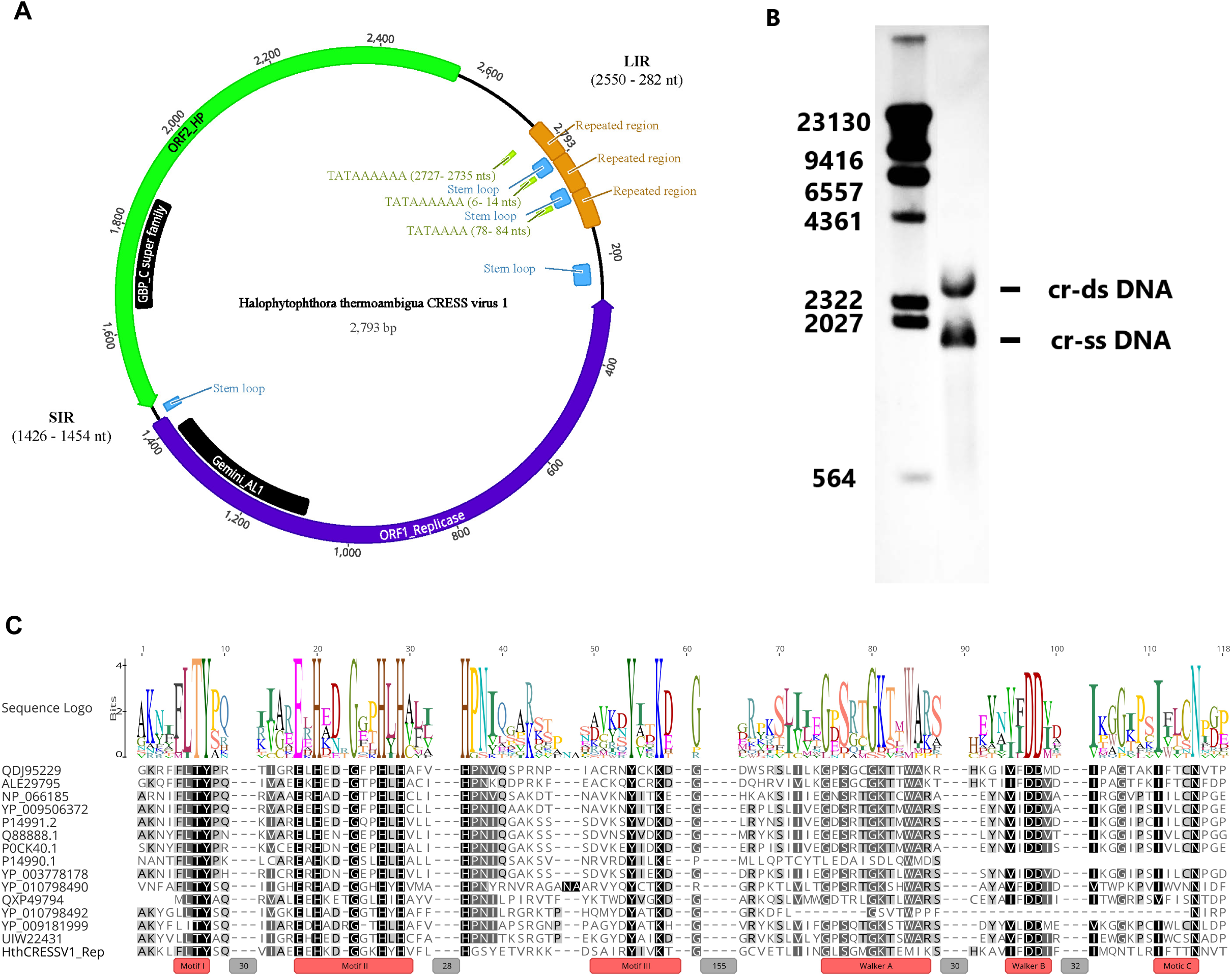
Genome organization and Rep motif architecture of HthCRESSV1. **(A)** Schematic representation of the circular genome of HthCRESSV1 showing the two major ORFs encoding the replication-associated protein (Rep) and the hypothetical protein (HP), as well as the large (LIR) and small (SIR) intergenic regions. Predicted stem–loop structures and repeated elements are indicated. **(B)** Southern blot analysis of HthCRESSV1. M, digoxigenin-labeled DNA size marker (λ DNA digested with HindIII). The viral DNA forms are indicated as follows: SC, supercoiled double-stranded (ds) DNA; cr, circular single-stranded (ss) DNA. **(C)** Comparison of Rep motif composition across representative members within the order *Geplafuvirales* and HthCRESSV1, including rolling-circle replication motifs (I–III) and helicase motifs (Walker A, Walker B and motif C). *Geplanaviridae:* QDJ95229 (Capybara virus 11), NP_066185 (Horseradish curly top virus); *Geminiviridae:* ALE29795 (Lake Sarah-associated circular virus 45); YP_009506372 (Abutilon golden mosaic Yucatan virus), P14991.2 (Beet curly top virus), Q88888.1 (Tomato pseudo-curly top virus), P0CK40.1 (Bean golden yellow mosaic virus), YP_003778178 (Turnip curly top virus); *Genomoviridae:* YP_010798490 (Zizania latifolia genomoviridae), YP_010798492 (Artemisia_carvifolia_genomoviridae), YP_009181999 (Soybean associated gemycircularvirus 1), UIW22431 (Sclerotinia sclerotiorum hypovirulence associated DNA virus 1), QXP49794 (Botrytis gemydayirivirus 2). Numbers in grey boxes represent number of nucleotides deleted in the alignment.

To further confirm the presence of HtCRESSV1 in *H. thermoambigua* and to assess the existence of the single-stranded and double-stranded replicative intermediates expected of viruses that replicate by rolling circle mechanism (as suggested by the presence of the *rep* gene), purified DNA samples obtained from mycelia were subjected to Southern hybridization with HtCRESSV1-specific probes. Two HtCRESSV1-specific bands were detected: one displayed faster mobility than the 2,027 bp marker of λDNA digested with HindIII, whereas the second one displayed slower mobility and exceeded 2,322 bp (Figures 1B). The distinct nature and size of the faster-migrating band are consistent with a circular ssDNA genome of approximately 2 kb, whereas the slower-migrating band is compatible with a dsDNA replicative intermediate. The presence of two defined viral DNA forms, rather than a heterogeneous smear indicative of degradation, is consistent with the replication of the viral genome as an episomal, likely circular molecule.

### Description of the genomic features of HthCRESSV1

The complete HtCRESSV1 genome contains two open reading frames (ORFs), both oriented in the same direction (Figure 1A). ORF1 encodes the putative Rep of 380 aa (estimated 44.66 kDa), with the domain organization typical of members in the phylum *Cressdnaviricota* [**5**], whereas ORF2 encodes a hypothetical protein (HP) of 363 aa (estimated 43.19 kDa). The two ORFs are separated by two intergenic regions. The smaller intergenic region (SIR) comprises a 29-nt AT-rich sequence located between the stop codon of the HP and the start codon of the Rep, and has the potential to form a stable secondary structure (Figure 1A and S2). The larger intergenic region (LIR) of 529 nt separates the 3’ end of Rep and the 5’ end of HP. A putative TATA-like promoter motif (TATATATATTACTA) is present at the 5′ end of the LIR, suggesting it may function as the transcriptional start site for the downstream HP gene. Notably, LIR features several tandemly repeated 72-nt units (TAGATTATAAAAAACCACAAGAAAGAATTGGTTTTTAGTTAAAAAGTTGGTTTTTAGGTTTTTTATATGGTT), encompassing fragments capable of forming stable stem-loop structures (Figures 1A and S2). The 3’ end of LIR includes a predicted stable hairpin with a 9-nucleotide loop, which could serve as the origin of replication, where the viral Rep initiates the rolling circle replication by introducing a nick. Notably, the sequence of the putative nonanucleotide, TGGCAGTGT, present at the apex of the hairpin is different from those predicted for other well studied cressdnaviricots, such as geminiviruses, genomoviruses, smacoviruses or circoviruses [**60–63**]. Thus, the overall architecture of the LIR, including its AT-richness, promoter-like motifs, and stem-loop structure support its potential dual role in initiating both replication and transcription.

### Comparative analysis of Rep and HP sequences

Multiple Rep sequence alignment demonstrated conservation of canonical rolling-circle replication (RCR) motifs I–III and helicase-associated Walker A, Walker B, and Motif C regions, confirming that HthCRESSV1 encodes a functional rolling-circle Rep protein (Figure 1C).

Maximum likelihood phylogenetic analysis placed HthCRESSV1 within a clade of unclassified viruses assigned to the order *Respensiviricetes*. This assemblage of unclassified viruses is more closely related to the family *Geplanaviridae* than to *Genomoviridae* or *Geminiviridae* (Figure 2A), highlighting its distinct evolutionary position within *Cressdnaviricota*. In addition to viral homologs, BLAST searches identified Rep-like sequences embedded in the genomes of several oomycetes, including *Lagenidium giganteum*, *Phytophthora sojae*, and *Peronospora tabacina* (Table S3). These sequences cluster with HthCRESSV1 in the phylogenetic tree, indicating shared evolutionary ancestry between the viral Rep and mitochondrial Rep-like elements in oomycetes.

**Figure 2.**
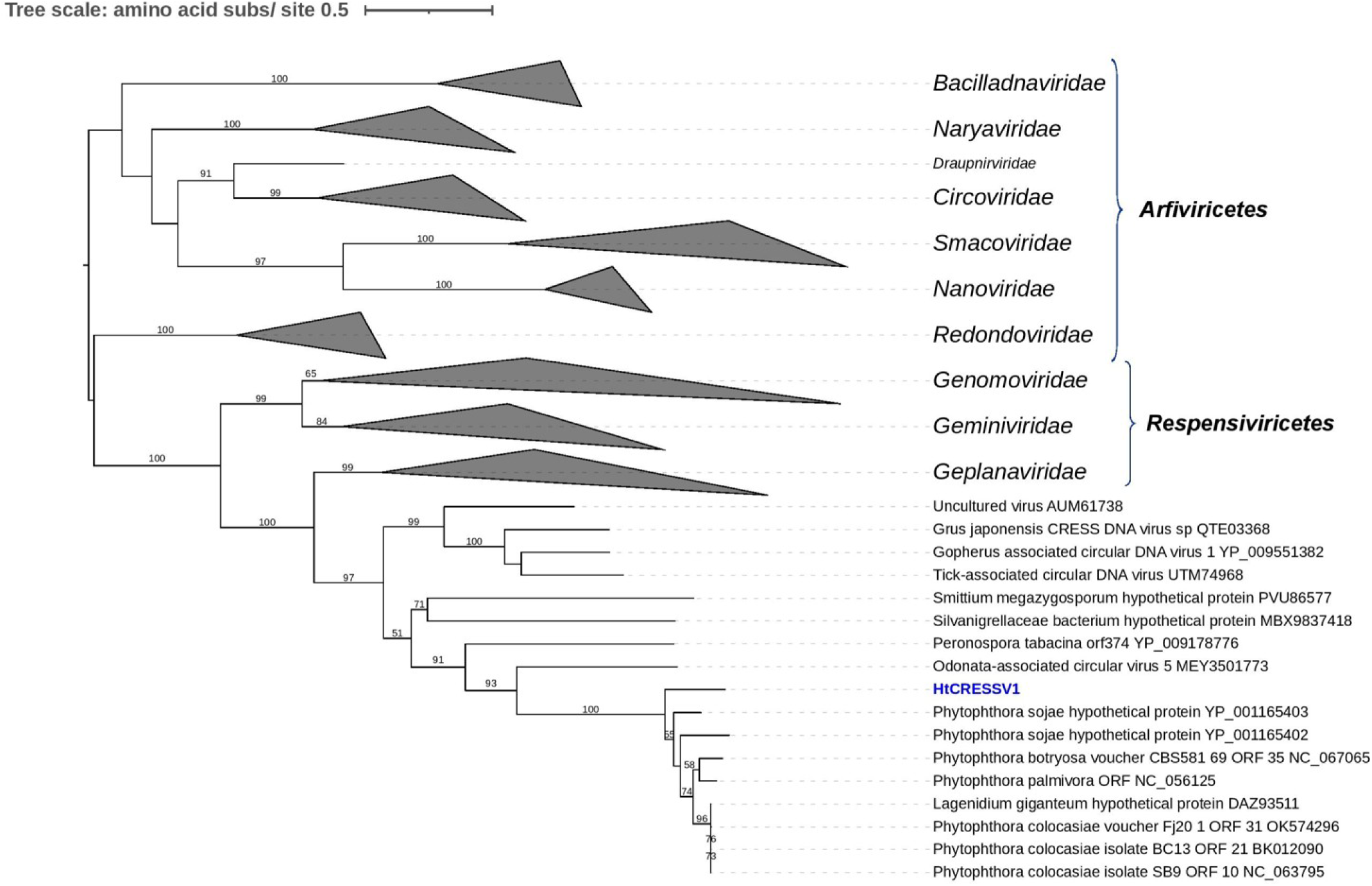
Randomized Axelerated ML tree (RAxML) depicting the phylogenetic relationships of the predicted Replicase **(A)** and the hypothetical protein **(B)** of HthCRESSV1 with their homologous found in oomycetes and other members of the phylum *Cressdnaviricota*. Nodes are labelled with bootstrap support values ≥50%. Branch lengths are scaled to the expected underlying number of substitutions per site. Some of the classified families and genera are collapsed to simplify the tree. Family classification and the corresponding GenBank accession numbers are shown next to the virus names. Scale bars represent expected changes per site per branch.

The BLASTn search revealed high nucleotide identity (76.88%) between HtCRESSV1 and an unannotated region of the mitochondrial genome of *Phytophthora palmivora* (culture ICMP:17709; NC_056125.1; 80% query coverage, E-value=0.0). Similar matches were detected in mitochondrial sequences of other oomycetes, including *Phytophthora colocasiae*, *Phytophthora botryosa*, and *Pseudoperonospora cubensis* (Table S4). These homologous regions included the coding regions of both HtCRESSV1 Rep (Table S4) and HP (Table S5). Alignment of the HtCRESSV1 HP with the homologous mitochondrial sequences revealed strong aa conservation across several regions (Figure S4).

In all characterized cressdnaviricots, the second of the two major ORFs encodes a capsid protein with the jelly-roll fold. Unexpectedly, sequence analysis showed that HP encoded by ORF2 does not have the potential to adopt the jelly-roll fold and instead is likely to be a membrane protein. The protein is predicted to be predominantly α-helical (70.2%), with β-stranded regions and unstructured coils contributing 6.6% and 23.2%, respectively. Five transmembrane domains were predicted in the N-terminus of the protein, whereas ∼50 C-terminus proximal amino acids were predicted to form a coiled-coil (Figure 3A). These features are unprecedented among proteins encoded by cressdnaviricots. Indeed, BlastP searches queried with the HP sequence against viral databases at NCBI and IMG/VR yielded no significant hits. Homologs of HP were exclusively encoded within the above-mentioned putative EVEs identified in oomycete mitochondria, suggesting specific association with HthCRESSV1-like viruses. To gain further insight into the topology and structure of HP, we attempted to model its structure using AlphaFold3 [51]. Given that AlphFold3 does not explicitly model membrane proteins in their natural membrane environment, we opted for modeling only the HP ectodomain first. To determine the likely oligomeric state of HP, we modeled multimers with progressively increasing oligomeric states from monomer to decamer and compared their predicted template modelling (pTM) and interface predicted template modelling (ipTM) scores (Table S6). The pTM and ipTM scores gradually improved from monomer to heptamer (best scores: ipTM=0.77, pTM=0.77), and started to decrease again for higher oligomeric states, suggesting that HP forms heptamers (Figures 3B, S1). In the modeled heptamer structure, the subunits arranged into an elongated tube-like structure, where the central α-helix formed contacts with the neighboring subunits. The predicted coiled-coil region was predicted to fold on itself, forming intramolecular contacts at the distal part of the ectodomain. We next modeled oligomers of the full-length HP. As in the case of the ectodomain, the highest pTM and ipTM scores were obtained for the hexameric and heptameric states, although the overall scores were considerably lower than for the ectodomain (Table S6). Nonetheless, the ectodomain model could be superposed with the full-length model (RMSD of 2 Å over 132 residues), supporting the relevance of the full-length heptameric model (Figures 3C, D). Thus, structural modeling suggests that HP forms an extended, membrane embedded spike-like complex. Structure-based searches with the monomer or heptamer against structure databases using DALI [**64**] and FoldSeek [**65**] did not reveal structural homologs, further emphasizing the distinctiveness of the HP compared to the previously characterized cellular and viral proteins.

**Figure 3.**
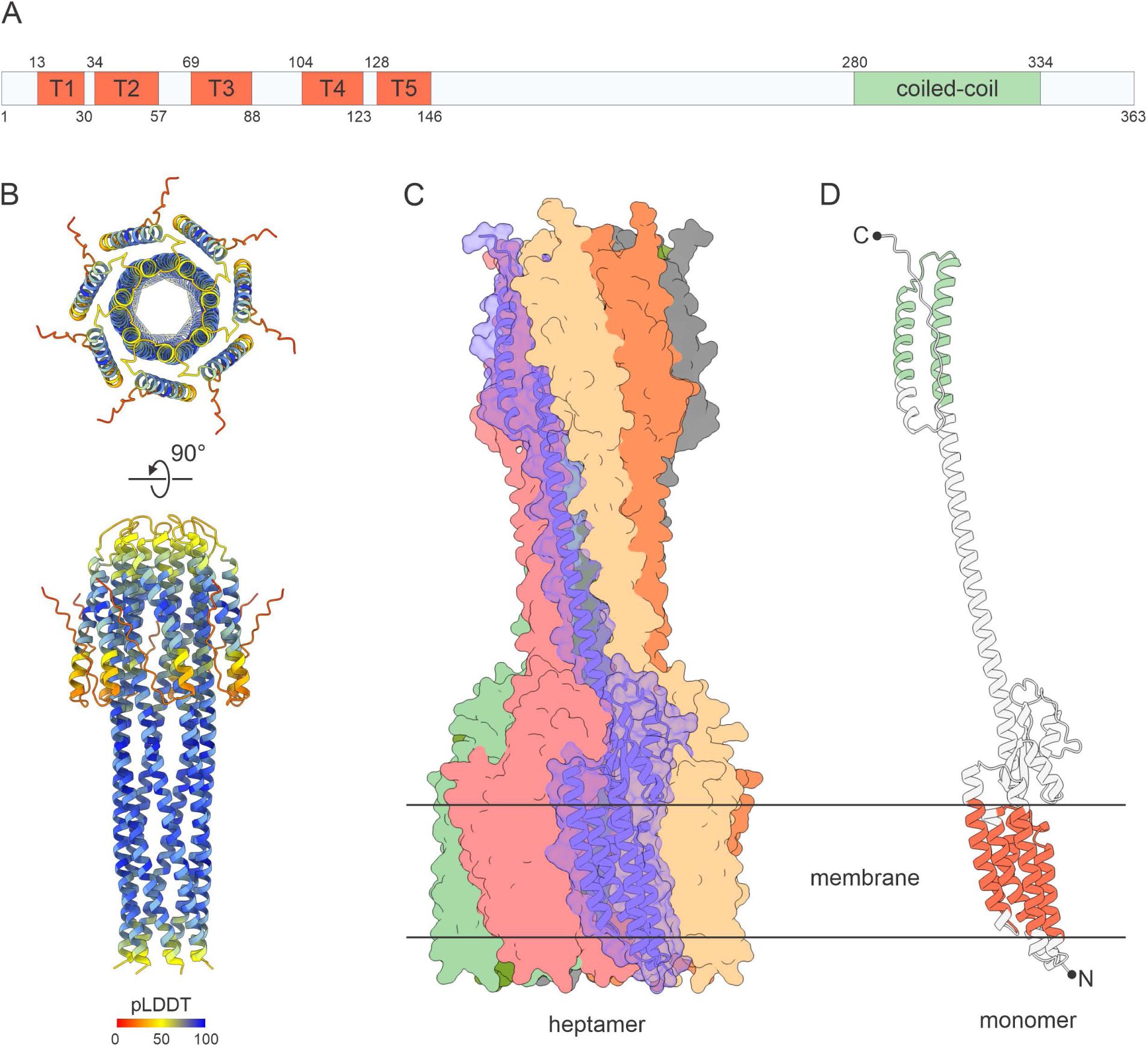
Structural modelling of the ORF2-encoded hypothetical protein. **(A)** Sequence features of the HthCRESSV1 hypothetical protein (HP). The sequence is represented by a rectangle, with the positions of the five transmembrane domains (T1-5) and the coiled-coil region indicated. **(B)** Structural model of the heptamer of the HP ectodomain generated with AlphaFold3. The model is colored based on the predicted local distance difference test (pLDDT) score, a per-residue measure of local confidence, from red to blue (the color scale is provided at the bottom of the figure). **(C)** Structural model of a heptamer of the full-length HP. Surface representation is shown, with each subunit colored differently. For one subunit, the surface is set to be semitransparent to show the secondary structure elements. **(D)** A monomer of the full-length HP. One of the subunits modeled in the context of a heptamer is shown separately and colored to depict the membrane-spanning domains and the coiled-coil region, as indicated in panel (A).

### Visualization of virus-like particles

The presence of viral particles has been demonstrated for a number of fungal cressdnaviricots. To verify whether HthCRESSV1 is capable of forming virus-like particles (VLP), despite lacking the gene for canonical capsid protein, we attempted to purify viral particles using standard viral particle purification protocols and visualised them by transmission electron microscopy (TEM; Figure 4A). Following fractionation of the putative VLP preparation on the continuous sucrose density gradient, 15 fractions were collected and analyzed for the presence of HthCRESSV1 genome using PCR with specific primers. HthCRESSV1 genomic DNA was abundant in fractions 4 and 5 (15–20% sucrose) (Figure 4B), suggesting the presence of HthCRESSV1 VLPs. TEM observation of these fractions revealed numerous spherical, pleomorphic particles. Consistent with the lack of the canonical capsid protein gene in the HthCRESSV1 genome, no icosahedral particles characteristic of previously characterized cressdnaviricots were observed (Figure 4C).

**Figure 4.**
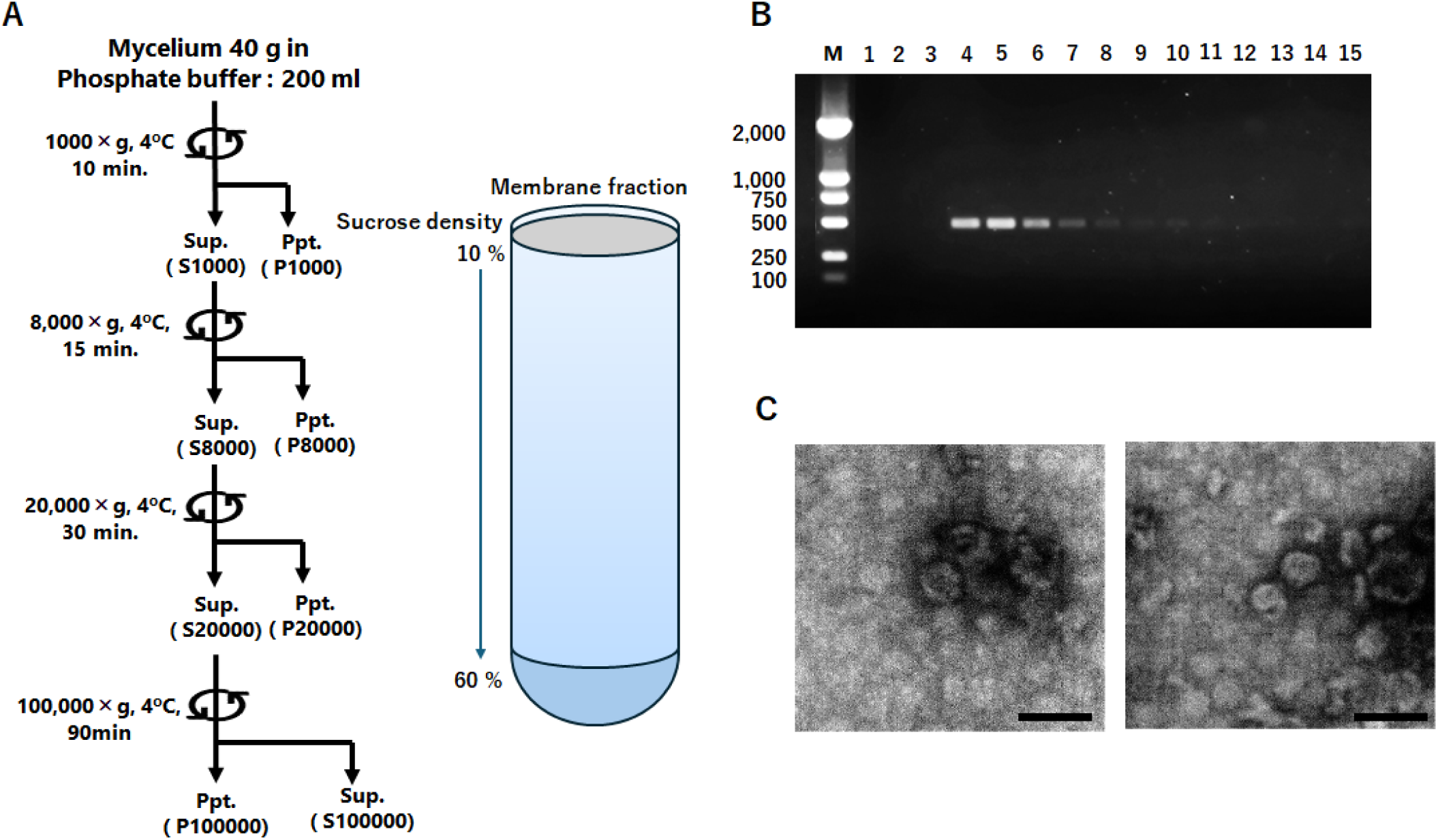
Purification attempts of HthCRESSV1 virus-like particles (VLPs). **(A)** Overview of VLP purification from strain BD651. Left: Mycelium homogenized in liquid nitrogen was suspended in isotonic solution and subjected to differential centrifugation at 1,000 *g*, 8,000 *g*, 20,000 *g*, and 100,000 *g*. Pellets (Ppt) and supernatants (Sup) were collected at each step and analyzed. Right: The P100,000 pellet was subjected to continuous sucrose density gradient centrifugation (10–60% w/v). Following ultracentrifugation, 2-ml aliquots were collected from the top, yielding 15 fractions. **(B)** PCR detection of HthCRESSV1 in the 15 fractions obtained by density gradient centrifugation. M, DNA size marker. **(C)** Transmission electron microscopy (TEM) of pooled fractions 4 and 5. Samples were negatively stained with 1% uranyl acetate. Scale bars, 50 nm.

### Analysis of HtCRESSV1 mitochondrial localization

Exclusive association of HtCRESSV1-like sequences with oomycete mitochondria prompted us to verify the mitochondrial localization of HtCRESSV1. To this end, the crude mitochondrial fraction (P12,000) obtained by cellular fractionation was subjected to two-step sucrose density gradient centrifugation. Following centrifugation, 15 fractions (F1-F15) containing purified mitochondria were collected (Figure 5A), total nucleic acids were extracted from each fraction and analyzed by PCR with HtCRESSV1-specific primers.

**Figure 5.**
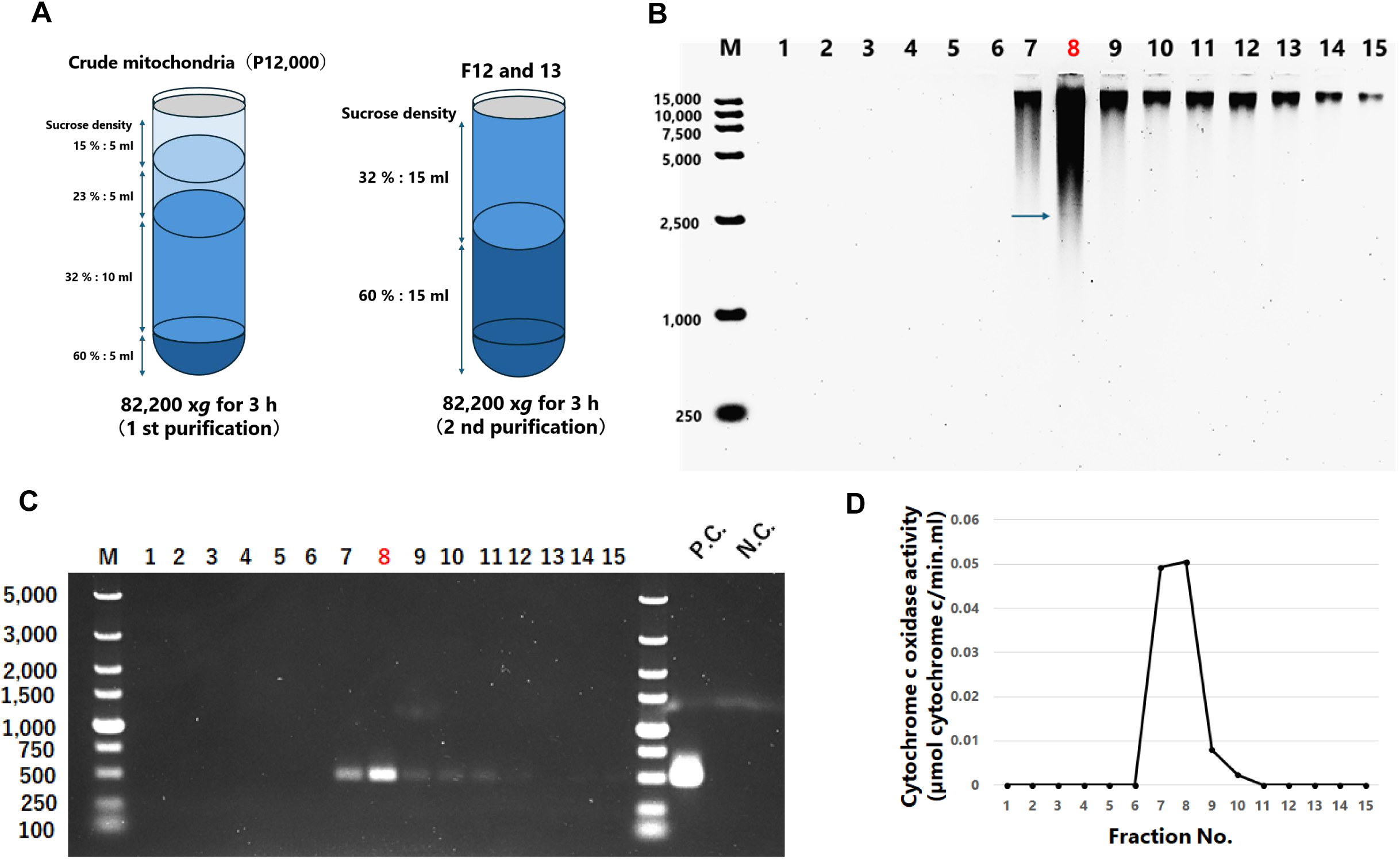
Mitochondrial localization of HthCRESSV1. **(A)** The P12,000 pellet was subjected to discontinuous sucrose density gradient centrifugation (15%, 23%, 32%, and 60% w/v). For the first purification, fractions collected at the 32%/60% interface (approximately fractions 12–13 from the top) were pooled and subjected to a second round of purification. Fractions (2 ml each) were collected from the top of the gradient. Total nucleic acids were extracted, ethanol-precipitated, and resuspended in TE buffer containing 1 mg ml⁻¹ RNaseA. **(B)** Agarose gel electrophoresis of total nucleic acids from the second mitochondrial purification fractions. Total nucleic acid samples extracted from each fraction were subjected to agarose gel electrophoresis from left (lane 1; 10%) to right (lane 15; 60%). The arrow indicates the HtCRESSV1 band. The image is shown with inverted contrast. M, DNA ladder (C) and (D) PCR specific for HthCRESSV1 detection and measurement of Cytochrome c oxidase (COX) activity for the second mitochondrial purification fractions. **(C)** HthCRESSV1-specific PCR of total nucleic acid samples extracted from each fraction. M, DNA ladder, P.C., positive control; N.C., negative control (water). **(D)** Cytochrome c oxidase (COX) activity in fractions from the second purification. COX activity was measured using 100 µl from each fraction, combined with 950 µl of assay buffer and 50 µl of substrate solution (reduced cytochrome c) for a total volume of 1,100 µl. Absorbance at 550 nm was measured for 1 minute, and the decrease in absorbance (ΔA) was recorded. Activity was calculated in units of µmol cytochrome c/min/ml using ΔA/min and the molar extinction coefficient difference of cytochrome c (21.8 mM⁻¹ cm⁻¹).

Agarose gel electrophoresis analysis of the total nucleic acids extracted from the 15 fractions revealed that fraction F8 contained the highest concentration of high molecular weight DNA, likely corresponding to the mitochondrial DNA. Notably, this fraction also contained a faint band of ∼2,500 bp in size, matching the expected size of the viral genome (Figure 5B). Consistently, semi-quantitative PCR analysis showed that HtCRESSV1 genome was enriched in fraction F8 (Figure 5C).

To evaluate the distribution of mitochondria in the gradient fractions, cytochrome oxidase (COX) activity, a mitochondrial marker, was measured in all 15 fractions. Strong COX activity was detected in fractions F7 and F8 (0.05 μmol cytochrome c/min/ml) indicating the presence of intact mitochondria at high concentrations in these fractions (Figure 5D), what strongly suggests that HtCRESSV1 localizes within mitochondria.

### HtCRESSV1 is transmitted via zoospores

To verify whether HtCRESSV1 can be also transmitted via zoospores we analyzed single-lesions formed by zoospore-derived isolates. All 25 strains derived from zoospores isolated from single lesions formed on cork oak leaves retained HtCRESSV1 infection (Figure 6A), indicating 100% zoospore transmission rate of the virus.

**Figure 6.**
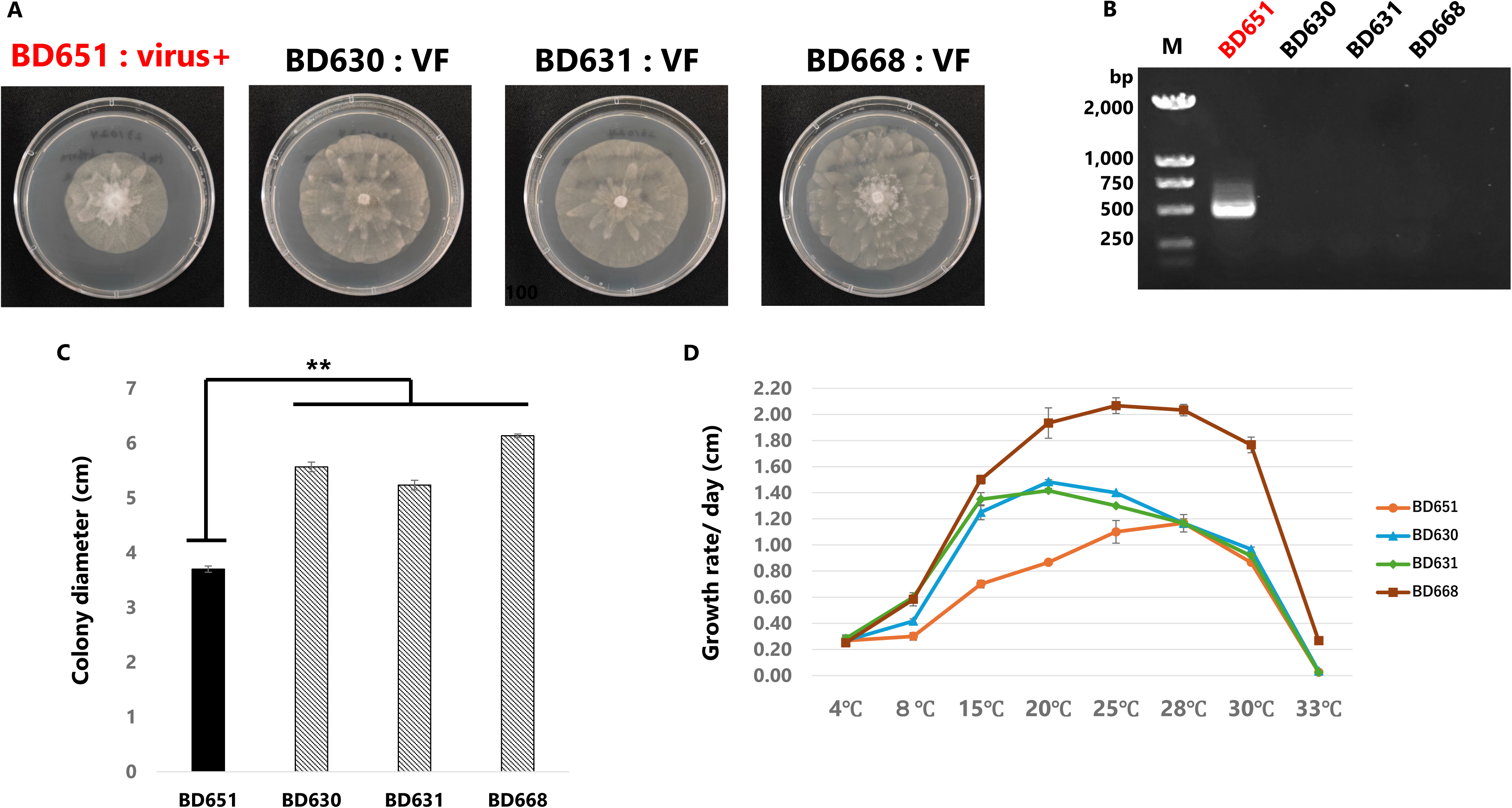
Biological effects and vertical transmission of HthCRESSV1. **(A–C)** Comparison of growth between virus-infected (BD651) and virus-free *H. thermoambigua* strains (BD630, BD631, BD668). **(A)** Colonies grown on s. V8A medium for 3 days. From left to right: BD651 (virus-infected), BD630 (virus-free), BD631 (virus-free), BD668 (virus-free). **(B)** PCR detection of HthCRESSV1 in each strain. M, DNA size marker. **(C)** Colony diameters after 3 days of growth on s. V8A medium. Black bar represents the virus-infected strain BD651; hatched bars represent virus-free strains. The virus-infected strain showed significantly reduced growth compared to virus-free strains (\**P* < 0.01). (D) Comparison of growth rates of BD651, BD630, BD631, and BD668 at different temperatures (4°C, 8°C, 15°C, 20°C, 25°C, 28°C, 30°C, and 33°C). Growth rates were calculated as colony diameter per day on s. V8A medium. Symbols: circles, BD651 (virus-infected); triangles, BD630 (virus-free); diamonds, BD631 (virus-free); squares, BD668 (virus-free). Note that the optimal growth temperature was 28°C for BD651, 20°C for BD630 and BD631, and 25°C for BD668.

To obtain virus-free strains, treatment with the antiviral agents, ribavirin and cycloheximide, a method frequently employed for mycovirus clearance [**59**], was performed. However, no virus-free strains were successfully isolated among the 20 oomycete strains regenerated from protoplasts (Figure S5). Similarly, the attempt to obtain virus-free strains by single-zoospore isolation was also unsuccessful, an approach used to cure fungal hosts from cressdnaviricot Diaporthe sojae circular DNA virus 1 [**19**].

### Evaluation of host impact

To investigate whether HtCRESSV1 infection affects the growth of the host oomycete, similar to what has been reported during the SsHDV1 infection of the fungus *Sclerotinia sclerotiorum*, we compared the growth of virus-infected strain BD651 and non-infected strains BD630, BD631 and BD668. First, mycelial colonies were compared after 3 days of incubation at 25°C on sV8A (prepared with 50% seawater). The colony diameter of the virus-infected strain BD651 was significantly smaller compared to the virus-free strains. The virus-infected strain also showed slightly higher hyphal density consistent with slower growth (Figures 6B, C).

Next, we compared the daily hyphal extension rates (cm/day) at temperatures ranging from 4°C to 33°C. The optimal growth temperatures (temperatures with the highest hyphal extension rates) for BD651, BD630, BD631 and BD668 were 28°C (1.17 cm/day), 20°C (1.48 cm/day), 20°C (1.42 cm/day), and 25°C (2.07 cm/day), respectively (Figures 6D). Thus, virus-infected strain BD651 had the highest optimal growth temperature, suggesting that HtCRESSV1 infection contributes to the host thermotolerance. Notably, however, the virus-infected strain BD651 was characterized by slower growth rate compared to the non-infected strains at lower (suboptimal) temperatures in the range of 4-25°C. These results suggest that HtCRESSV1 infection negatively impacts growth rates of the oomycete across a wide temperature range, most likely reducing its ability to colonise plant tissues.

## DISCUSSION

In this study, we identified a novel virus with circular ssDNA genome, HthCRESSV1, associated with the marine oomycete *Halophytophthora thermoambigua*. Although viruses with linear ssDNA genomes classified in the recently created family *Oomyviridae* (phylum *Cressdnaviricota*, class *Arfiviricetes*, order *Lineavirales*) have been predicted to infect oomycetes based on the presence of related EVEs in the genomes of diverse oomycetes species [**20**], to the best of our knowledge, HthCRESSV1 represents the first experimentally characterized cressdnaviricot infecting an oomycete. Indeed, all mycoviruses infecting marine fungi or oomycetes isolated up to now have RNA genomes [**30–33**]. This is in line with the fact that most known mycoviruses, regardless of the ecosystem they inhabit, have dsRNA or ssRNA genomes, whereas ssDNA mycoviruses are relatively rare [**27**]. Notably, HthCRESSV1 is unrelated to *Oomyviridae* [**20**], which have linear genomes, encode a typical jelly-roll capsid protein and based on the Rep phylogeny are placed within the class *Arfiviricetes*. By contrast, Rep phylogeny firmly places HthCRESSV1-like viruses within the class *Repensiviricetes*, as a separate family, which we propose naming ‘Mitodnaviridae’.

The only other characterized group of ssDNA viruses infecting stramenopile hosts is the family *Bacilladnaviridae*, which, like *Oomyviridae,* is clasisfied within the class *Arfiviricetes*. Bacilladnaviruses infect marine diatoms and induce host cell lysis, releasing progeny virions into the surrounding seawater [**66, 67**]. Thus, they are transmitted horizontally in marine ecosystems without the involvement of specific vectors. Whether HthCRESSV1 is capable of horizontal transmission between susceptible hosts remains unknown. It also remains unclear whether observed pleomorphic HthCRESSV1-associated particles contribute to this process and to what extent they provide protection to the viral genome outside the host in the marine environment.

The repetitive structures found in the HthCRESSV1 LIR resemble the abundant tandem and palindromic repeats that characterize mitochondrial genomes and are involved in recombination, genome rearrangement, and control-region function [**68–72**]. This similarity raises the possibility that HthCRESSV1 originated from an endogenized mitochondrial episome or mobile viral element that gained autonomy while retaining compatibility with mitochondrial sequence environments. Such a model aligns with current views of cressdnaviricots’ evolution, which suggest recurrent recombination among plasmid-like replicons and repeated acquisition or loss of accessory genes, including capsid proteins, throughout eukaryotic evolution [**73–76**]. In this framework, the Gemini_AL1-like nuclease–helicase domain in HthCRESSV1 Rep protein reflects conserved ancestry within *Repensiviricetes* rather than indicating a specific relationship with geminiviruses. The absence of any recognizable structural or capsid-associated protein suggests that HthCRESSV1 may represent a lineage in which the canonical jelly-roll capsid has been secondarily lost or replaced. Within the context of the proposed ‘Mitodnaviridae’, this feature is consistent with an intracellular replication strategy associated with mitochondrial localization, where selective constraints on extracellular transmission may be reduced. Comparable processes, particularly modular genome exchange and structural gene turnover, have been widely documented in CRESS-DNA viruses and related elements, linking plasmid-like replicons and virion-forming viruses [**3–5, 75, 76**].

From a virological perspective, most DNA viruses replicate in the nucleus, and it remains unclear whether minimal cressdnaviricot viruses, encoding only Rep and Cap, possess the capacity to actively cross the mitochondrial double membrane. Information on viruses that directly interact with mtDNA is extremely limited. For example, mitoviruses, which replicate within the mitochondrial matrix, have left mitovirus-like sequences in both the nuclear and mitochondrial genomes of several vascular plants [**77–79**]. Among infectious DNA viruses, hepatitis B virus (HBV) not only integrates into the host nuclear genome but has also been shown to deliver HBV RNA into mitochondria of hepatocellular carcinoma cells, where it may participate in mtDNA integration [**80**]. This process is mediated by polynucleotide phosphorylase (PNPASE), which facilitates RNA transport into mitochondria. In contrast, our findings indicate that the DNA genome of HthCRESSV1 is present within mitochondria, and given the RNA specificity of PNPASE, its translocation likely involves a distinct mechanism. Further studies using immunoelectron microscopy or other imaging approaches will be required to determine whether HthCRESSV1 localizes to the mitochondrial matrix or the intermembrane space.

The ORF2 of HthCRESSV1 is predicted to encode an α-helical membrane protein with five N-terminal transmembrane domains and a C-terminal coiled-coil region. Structural modeling suggests that this protein can form multimeric complexes within membranes, likely as trimers, based on its structural similarity to phage tail spike proteins. Although the confidence in the transmembrane topology prediction is moderate, the extracellular (ectodomain) structure is well supported. Homologs of ORF2 were not detected in known members of the phylum *Cressdnaviricota* but were identified in several oomycete mitochondrial genomes [**81–85**], suggesting an evolutionary relationship with mobile mitochondrial elements or integrated viral fragments. These homologs often occur without adjacent *Rep* genes, implying independent acquisition through horizontal transfer or secondary loss of *Rep*. The tight association of the HthCRESSV1 genome with mitochondrial DNA, coupled with the absence of nuclear integration, supports the hypothesis that replication occurs within mitochondria. This is further reinforced by TEM, which revealed vesicle-like structures potentially associated with viral replication. Functionally, ORF2 likely contributes to membrane dynamics. If the viral genome is encapsulated within membrane-bound vesicles, ORF2 may act as a fusogen, mediating vesicle formation, budding, or membrane fusion events that facilitate replication, genome packaging, or evasion of mitochondrial antiviral defenses [**86, 87**]. Alternatively, if the virus remains confined within the mitochondrial compartment, ORF2 may remodel inner membrane structures such as cristae or mitovesicles to create specialized niches for replication [**88, 89**]. The presence of a C-terminal coiled-coil domain supports a role in oligomerization or scaffolding, common features of membrane-associated structural or assembly proteins [**90, 91**]. Given its distinctive combination of features and phylogenetically isolated position, ORF2 likely represents a novel membrane-active viral factor, possibly acquired through horizontal transfer from a mitochondrial mobile element or host organellar DNA.

Consistent with this proposed mitochondrial association, indirect evidence supporting the interaction between HthCRESSV1 and mitochondria includes its highly efficient vertical transmission via zoospores and variation in host growth across temperatures. The species epithet of *H. thermoambigua* reflects its ambiguous optimal growth temperature [**38**]; comparative growth assays showed that, compared to the virus-free strains, the virus-infected strain BD651 exhibited a reduced growth overall while maintaining relatively similar growth at higher temperatures. Mitochondrial genomes have been implicated in thermal adaptation in several organisms, including *Phytophthora infestans*, *Drosophila melanogaster*, and *Saccharomyces cerevisiae* [**71, 92, 93**]. Although HthCRESSV1 is not integrated into mtDNA, its association with mitochondria may influence organelle function and contribute to reduce growth rates and enhance optimal growth temperature. In this context, mitochondrial RNA viruses, mitoviruses, provide a useful comparison. Although phylogenetically unrelated to HthCRESSV1 and typically associated with cryptic infections, they share a common intracellular niche and are transmitted vertically through spores very efficiently [**94**]. In some cases, some have been linked to altered host phenotypes; for example, infection by Sclerotinia sclerotiorum mitovirus 1 (SsMV1) in *Sclerotinia sclerotiorum* is associated with reduced growth and hypovirulence, alongside mitochondrial abnormalities [**78**].

In this study, attempts to eliminate HthCRESSV1 using ribavirin were unsuccessful, and the virus remained stably transmitted through zoospores. Similar persistence has been reported for other cressdnaviricots infecting fungi [**19**]. Notably, successful re-infection has been demonstrated for certain viruses such as SsHADV1 using purified virions or infectious clones [**15, 19**]. Developing a re-infection system for HthCRESSV1, potentially through mitochondrial transfection approaches, will be essential to clarify its biological effects on the host.

## Conclusion

HthCRESSV1 represents a circular DNA virus with unique genomic features and a close association with oomycete mitochondria. Its genome encodes a divergent Rep protein and a membrane-associated HP conserved in multiple mitogenomes, suggesting prolonged interaction with the mitochondrial compartment. The absence of recognizable capsid genes and the presence of repetitive regulatory elements point to a separated evolutionary trajectory within cressdnaviricots. Together, these findings expand current views of CRESS virus diversity and highlight mitochondria as a potential niche for circular DNA replicons in eukaryotes.

## Author Contributions

Conceptualization, L.B. and K.S.; methodology, K.S., L.B., C.M.; formal analysis, O.H., K.S., L.B., M.K.; writing—original draft preparation, K.S., L.B.; writing: K.S., L.B., M.K., M.H.J, O.H. and T.J.; funding acquisition: T.J., H.M., M.H.J., L.B., K.S. All authors have read and agreed to the published version of the manuscript.

---

**Supplementary Figure S1. Genome validation and assembly of HthCRESSV1**

**(A)** PCR amplification of overlapping genomic fragments using different primer combinations. **(B)** Schematic representation of primer binding sites on the original contig. **(C)** Coverage plots showing amplicon-derived sequences mapped to the HthCRESSV1 genome in Geneious Prime. **(D)** Read coverage and orientation across the finalized genome assembly.

**Supplementary Figure S2. Predicted stem–loop structures in selected genomic regions,** including the SIR **(A)** and internal genomic fragments **(B–C)**, generated using minimum free energy folding models. Free energy values, ensemble diversity, and nucleotide positions are indicated.

**Supplementary Figure S3. Amino acid alignment showing conserved motifs shared by the hypothetical protein of HthCRESSV1 with mitochondrial putative proteins other related of oomycetes.**

**Supplementary Figure S4. Structural modelling of the oligomers of the HP ectodomain.** The figure shows multimeric states of the HP ectodomain from monomer to decamer colored based on the predicted local distance difference test (pLDDT) score, a per-residue measure of local confidence, from red to blue (the color scale is provided in the top right corner of the figure). The predicted template modelling (pTM) and the interface predicted template modelling (ipTM) scores are also indicated under each model.

**Supplementary Figure S5. Attempts to eliminate HthCRESSV1 from strain BD651. (A)** PCR detection of HthCRESSV1 in 25 strains obtained by single zoospore isolation. M, DNA size marker. **(B)** PCR detection of HthCRESSV1 in protoplast-regenerated strains exposed to ribavirin (300 µg ml⁻¹ final concentration) and cycloheximide (5 µg ml⁻¹ final concentration) in regeneration medium. Regenerated mycelia were subsequently transferred to s. V8A medium without ribavirin and cycloheximide. M, DNA size marker.

## References

1. Rosario, K., Duffy, S. & Breitbart, M. A field guide to eukaryotic circular single-stranded DNA viruses: Insights gained from metagenomics. Arch Virol 157 (2012).

2. Krupovic, M. Networks of evolutionary interactions underlying the polyphyletic origin of ssDNA viruses. Curr Opin Virol 3 (2013).

3. Zhao, L., Rosario, K., Breitbart, M. & Duffy, S. Eukaryotic circular rep-encoding single-stranded DNA (CRESS DNA) viruses: ubiquitous viruses with small genomes and a diverse host range. Adv Virus Res 103 (2019).

4. Krupovic, M., et al. *Cressdnaviricota*: a virus phylum unifying seven families of Rep-encoding viruses with single-stranded, circular DNA genomes. J Virol 94, (2020).

5. Kazlauskas, D., Varsani, A., Koonin, E. v. & Krupovic, M. Multiple origins of prokaryotic and eukaryotic single-stranded DNA viruses from bacterial and archaeal plasmids. Nat Commun 2019 10:1 10, 3425- (2019).

6. Kulshrestha, S., Bhardwaj, A. & Vanshika. Geminiviruses: Taxonomic structure and diversity in genomic organization. Recent Pat Biotechnol 14, 86–98 (2020).

7. Diemer, G. S. & Stedman, K. M. A novel virus genome discovered in an extreme environment suggests recombination between unrelated groups of RNA and DNA viruses. Biol Direct 7, (2012).

8. Roux, S. et al. Chimeric viruses blur the borders between the major groups of eukaryotic single-stranded DNA viruses. Nat Commun 4, (2013).

9. Kazlauskas, D. et al. Evolutionary history of ssDNA bacilladnaviruses features horizontal acquisition of the capsid gene from ssRNA nodaviruses. Virol 504, 114–121 (2017).

10. Munke, A. et al. Primordial capsid and spooled ssDNA genome structures unravel ancestral events of eukaryotic viruses. mBio 13, (2022).

11. Simmonds, P. et al. Changes to virus taxonomy, the international code of virus classification and nomenclature, and the ICTV statutes ratified by the International Committee on Taxonomy of Viruses (2025). Archi Virol, 171 (2025).

12. Krupovic, M., Dolja, V. v. & Koonin, E. v. The LUCA and its complex virome. Nat Rev Microbiol 18, (2020).

13. Varsani, A. & Krupovic, M. 2024 *Smacoviridae* family update: 59 new species in seven genera. Arch Virol 169, (2024).

14. Varsani, A. et al. 2024 taxonomy update for the family *Circoviridae*. Arch Virol 169, (2024).

15. Yu, X. et al. A geminivirus-related DNA mycovirus that confers hypovirulence to a plant pathogenic fungus. PNAS 107, 8387–92 (2010).

16. Li, P. et al. A tripartite ssDNA mycovirus from a plant pathogenic fungus is infectious as cloned DNA and purified virions. Sci Adv 6, (2020).

17. Hao, F., Wu, M. & Li, G. Characterization of a novel genomovirus in the phytopathogenic fungus *Botrytis cinerea*. Virol 553, (2021).

18. Ruiz-Padilla, A., Turina, M. & Ayllón, M. A. Molecular characterization of a tetra segmented ssDNA virus infecting *Botrytis cinerea* worldwide. Virol J 20, (2023).

19. Wang, X. et al. A circular single-stranded DNA mycovirus infects plants and confers broad-spectrum fungal resistance. Mol Plant 17, 955–971 (2024).

20. Sabanadzovic, S. et al. Summary of taxonomy changes ratified by the International Committee on Taxonomy of Viruses (ICTV) from the Fungal and Protist Viruses Subcommittee, 2025. J Gen Virol 106, (2025).

21. Eaglesham, J. B. & Hewson, I. Widespread detection of circular replication initiator protein (rep)-encoding ssDNA viral genomes in estuarine, coastal and open ocean net plankton. Mar Ecol Prog Ser 494, 65–72 (2013).

22. Reavy, B. et al. Distinct circular single-stranded DNA viruses exist in different soil types. Appl Environ Microbiol 81, 3934–3945 (2015).

23. Rosario, K. et al. Virus discovery in all three major lineages of terrestrial arthropods highlights the diversity of single-stranded DNA viruses associated with invertebrates. PeerJ 2018, e5761 (2018).

24. Fehér, E. et al. Genomic diversity of cress dna viruses in the eukaryotic virome of swine faeces. Microorganisms 9, (2021).

25. Varsani, A. & Krupovic, M. Family *Genomoviridae*: 2021 taxonomy update. Arch Virol 166, 2911–2926 (2021).

26. Zhao, L., Lavington, E. & Duffy, S. Truly ubiquitous CRESS DNA viruses scattered across the eukaryotic tree of life. J Evol Biol 34, 1901–1916 (2021).

27. Kondo, H., Botella, L. & Suzuki, N. Mycovirus Diversity and Evolution Revealed/Inferred from Recent Studies. Annu Rev Phytopathol 60, 307–336 (2022).

28. Suttle, C. A. Viruses in the sea. Nature 437, 356–361 (2005).

29. Coy, S. R., Gann, E. R., Pound, H. L., Short, S. M. & Wilhelm, S. W. Viruses of eukaryotic algae: Diversity, methods for detection, and future directions. Viruses 10 (2018).

30. Nerva, L. et al. Multiple approaches for the detection and characterization of viral and plasmid symbionts from a collection of marine fungi. Virus Res 219, 22–38 (2016).

31. Nerva, L. et al. The mycovirome of a fungal collection from the sea cucumber *Holothuria polii*. Virus Res 273, 197737 (2019).

32. Botella, L. et al. Marine oomycetes of the genus *Halophytophthora* harbor viruses related to bunyaviruses. Front Microbiol 11, 1–13 (2020).

33. Botella, L. & Jung, T. Multiple viral infections detected in *Phytophthora condilina* by total and small RNA sequencing. Viruses 13, 620 (2021).

34. Sullivan, B. K., Trevathan-Tackett, S. M., Neuhauser, S. & Govers, L. L. Review: Host-pathogen dynamics of seagrass diseases under future global change. Mar Pollut Bull 134, 75–88 (2018).

35. Govers, L. L. et al. Marine *Phytophthora* species can hamper conservation and restoration of vegetated coastal ecosystems. Proc. Biol. Sci 283 (2016).

36. Newell, S. Y. & Fell, J. W. Distribution and experimental responses to substrate of marine oomycetes (*Halophytophthora* spp.) in mangrove ecosystems. Mycol Res 96, 851–856 (1992).

37. Newell, S. Y. & Fell, J. W. Competition among mangrove oomycotes, and between oomycotes and other microbes. Aquat Microb Ecol 12, 21–28 (1997).

38. Maia, C. et al. Eight new *Halophytophthora* species from marine and brackish-water ecosystems in Portugal and an updated phylogeny for the genus. Persoonia 48, 54–90 (2022).

39. Jung, T. et al. Diversity of *Phytophthora* species in natural ecosystems of Taiwan and association with disease symptoms. Plant Pathol 66, 194–211 (2017).

40. Chomczynski, P., Wilfinger, W., Kennedy, A., Rymaszewski, M. & Mackey, K. RNAzol® RT: a new single-step method for isolation of RNA. Nat Methods 7, 4–5 (2010).

41. Bushnell, B., Rood, J. & Singer, E. BBMerge – Accurate paired shotgun read merging via overlap. PLoS One 12, (2017).

42. Martin, M. Cutadapt removes adapter sequences from high-throughput sequencing reads. EMBnet J 17, (2011).

43. Kopylova, E., Noé, L. & Touzet, H. SortMeRNA: Fast and accurate filtering of ribosomal RNAs in metatranscriptomic data. Bioinformatics 28, (2012).

44. Dobin, A. STAR Manual. Github.Com (2014).

45. Bankevich, A. et al. SPAdes: A new genome assembly algorithm and its applications to single-cell sequencing. J Comput Biol 19, (2012).

46. Camacho, C. et al. BLAST+: Architecture and applications. BMC Bioinformatics 10, (2009).

47. Sims, D., Sudbery, I., Ilott, N. E., Heger, A. & Ponting, C. P. Sequencing depth and coverage: key considerations in genomic analyses. Nat Rev Genet 2014 15:2 15, 121–132 (2014).

48. Stamatakis, A. RAxML-VI-HPC: Maximum likelihood-based phylogenetic analyses with thousands of taxa and mixed models. Bioinformatics 22, 2688–2690 (2006).

49. Letunic, I., & Bork, P. Interactive Tree Of Life (iTOL) v5: an online tool for phylogenetic tree display and annotation. Nucleic Acids Res 49, W293–W296 (2021).

50. Buchan, D. W. A., Moffat, L., Lau, A., Kandathil, S. M. & Jones, D. T. Deep learning for the PSIPRED Protein Analysis Workbench. Nucleic Acids Res 52, W287–W293 (2024).

51. Zimmermann, L. et al. A Completely reimplemented MPI bioinformatics toolkit with a new HHpred server at its core. J Mol Biol 430, 2237–2243 (2018).

52. Abramson, J. et al. Accurate structure prediction of biomolecular interactions with AlphaFold 3. Nature 630, 493–500 (2024).

53. Meng, E. C. et al. UCSF ChimeraX: Tools for structure building and analysis. Protein Sci 32, (2023).

54. Sakuta, K., Uchida, K., Fukuhara, T., Komatsu, K., Okada, R., & Moriyama, H. Successful full-length genomic cloning and characterization of site-specific nick structures of Phytophthora endornaviruses 2 and 3 in yeast, Saccharomyces cerevisiae. Front Microbio 14, 1243068 (2023).

55. Okada, R., Kiyota, E., Moriyama, H., Fukuhara, T. & Natsuaki, T. A simple and rapid method to purify viral dsRNA from plant and fungal tissue. J Gen Plant Pathol 81, 103–107 (2015).

56. Jesus, A. L. et al. The genus *Halophytophthora* (Peronosporales, Straminipila) in Brazil: first descriptions of species. Brazilian J Bot 39, 729–739 (2016).

57. Jung, T., Cooke, D. E. L., Blaschke, H., Duncan, J. M. & Oßwald, W. *Phytophthora quercina* sp. nov., causing root rot of European oaks. Mycol Res 103, 785–798 (1999).

58. Sakuta, K. et al. Novel endornaviruses infecting *Phytophthora cactorum* that attenuate vegetative growth, promote sporangia formation and confer hypervirulence to the host oomycete. J Gen Virol 106, (2025).

59. Uchida, K. et al. Two novel Endornaviruses co-infecting a *Phytophthora* pathogen of *Asparagus officinalis* modulate the developmental stages and fungicide sensitivities of the host oomycete. Front Microbiol 12, (2021).

60. Varsani, A. & Krupovic, M. Sequence-based taxonomic framework for the classification of uncultured single-stranded DNA viruses of the family *Genomoviridae*. Virus Evol 3, (2017).

61. Varsani, A. & Krupovic, M. *Smacoviridae*: a new family of animal-associated single-stranded DNA viruses. Arch Virol 163, 2005–2015 (2018).

62. Fiallo-Olivé, E. et al. ICTV Virus Taxonomy Profile: *Geminiviridae* 2021. J Gen Virol 102, (2021).

63. Varsani, A. et al. 2024 taxonomy update for the family *Circoviridae*. Arch Virol 169, (2024).

64. Holm, L. Dali server: structural unification of protein families. Nucleic Acids Res 50, W210–W215 (2022).

65. van Kempen, M. et al. Fast and accurate protein structure search with Foldseek. Nat Biotechnol 42, 243–246 (2024).

66. Tomaru, Y., Shirai, Y., Suzuki, H., Nagumo, T. & Nagasaki, K. Isolation and characterization of a new single-stranded DNA virus infecting the cosmopolitan marine diatom *Chaetoceros debilis*. Aquat Microb Ecol 50, (2008).

67. Danovaro, R. et al. Major viral impact on the functioning of benthic deep-sea ecosystems. Nature 454, (2008).

68. Kroon, L. P. N. M., Bakker, F. T., van den Bosch, G. B. M., Bonants, P. J. M. & Flier, W. G. Phylogenetic analysis of *Phytophthora* species based on mitochondrial and nuclear DNA sequences. Fungal Genet Bio 41, 766–782 (2004).

69. Tangphatsornruang, S. et al. Comparative mitochondrial genome analysis of *Pythium insidiosum* and related oomycete species provides new insights into genetic variation and phylogenetic relationships. Gene 575, (2016).

70. Casabella-Herrero, G., Martínez-Ríos, M., Viljamaa-Dirks, S., Martín-Torrijos, L. & Diéguez-Uribeondo, J. Aphanomyces astaci mtDNA: insights into the pathogen’s differentiation and its genetic diversity from other closely related oomycetes. Fungal Biol 125, 316–325 (2021).

71. Shen, L. L. et al. Mitochondrial genome contributes to the thermal adaptation of the oomycete *Phytophthora infestans*. Front Microbiol 13, (2022).

72. Wang, J. et al. Plant organellar genomes: much done, much more to do. Trends Plant Sci 29, 754–769 (2024).

73. Kazlauskas, D., Varsani, A. & Krupovic, M. Pervasive chimerism in the replication-associated proteins of uncultured single-stranded DNA viruses. Viruses 10, (2018).

74. Desingu, P. A. & Nagarajan, K. Genetic diversity and characterization of circular replication (Rep)-encoding single-stranded (CRESS) DNA viruses. Microbiol Spectr 10, (2022).

75. Koonin, E. v., Dolja, V. v. & Krupovic, M. Origins and evolution of viruses of eukaryotes: The ultimate modularity. Virology 479–480, 2–25 (2015).

76. Krupovic, M. & Koonin, E. v. Multiple origins of viral capsid proteins from cellular ancestors. PNAS 114, E2401–E2410 (2017).

77. Bruenn, J. A., Warner, B. E. & Yerramsetty, P. Widespread mitovirus sequences in plant genomes. PeerJ 3, (2015).

78. Xu, Z. et al. A mitovirus related to plant mitochondrial gene confers hypovirulence on the phytopathogenic fungus *Sclerotinia sclerotiorum*. Virus Res 197, 127–136 (2014).

79. di Silvestre, D. et al. Presence of a mitovirus is associated with alteration of the mitochondrial proteome, as revealed by protein–protein interaction (PPI) and co-expression network models in *Chenopodium quinoa* plants. Biology 2022, Vol. 11, Page 95 11, 95 (2022).

80. Giosa, D. et al. Mitochondrial DNA is a target of HBV integration. Commun Biol 6, (2023).

81. Martin, F. N., Bensasson, D., Tyler, B. M. & Boore, J. L. Mitochondrial genome sequences and comparative genomics of *Phytophthora ramorum* and *P. sojae*. Curr Genet 51, 285–296 (2007).

82. Winkworth, R. C. et al. Comparative analyses of complete Peronosporaceae (Oomycota) mitogenome sequences-insights into structural evolution and phylogeny. Genome Biol Evol 14, (2022).

83. Winkworth, R. C. et al. A LAMP at the end of the tunnel: A rapid, field deployable assay for the kauri dieback pathogen, *Phytophthora agathidicida*. PLoS One 15, (2020).

84. Derevnina, L. et al. Genome Sequence and Architecture of the Tobacco Downy Mildew Pathogen Peronospora tabacina. Mol Plant Microbe Interact 28, 1198–1215 (2015).

85. Morgan, W. R. & Tartar, A. Decontamination and annotation of the draft genome sequence of the oomycete *Lagenidium giganteum* ARSEF 373. Microbiol Resour Announc 12, (2023).

86. Miller, S. & Krijnse-Locker, J. Modification of intracellular membrane structures for virus replication. Nat Rev Microbiol 2008 6:5 6, 363–374 (2008).

87. Hsu, N. Y. et al. Viral reorganization of the secretory pathway generates distinct organelles for RNA replication. Cell 141, 799–811 (2010).

88. Kopek, B. G., Perkins, G., Miller, D. J., Ellisman, M. H. & Ahlquist, P. Three-dimensional analysis of a viral RNA replication complex reveals a virus-induced mini-organelle. PLoS Biol 5, 2022–2034 (2007).

89. den Boon, J. A. & Ahlquist, P. Organelle-like membrane compartmentalization of positive-strand RNA virus replication factories. Annu Rev Microbiol 64, 241–256 (2010).

90. Burkhard, P., Stetefeld, J. & Strelkov, S. v. Coiled coils: A highly versatile protein folding motif. Trends Cell Biol 11, 82–88 (2001).

91. Lupas, A. N. & Bassler, J. Coiled Coils – A Model System for the 21st Century. Trends Biochem Sci 42, 130–140 (2017).

92. Camus, M. F., Wolff, J. N., Sgrò, C. M. & Dowling, D. K. Experimental support that natural selection has shaped the latitudinal distribution of mitochondrial haplotypes in Australian *Drosophila melanogaster*. Mol Biol Evol 34, 2600–2612 (2017).

93. Li, X. C., Peris, D., Hittinger, C. T., Sia, E. A. & Fay, J. C. Mitochondria-encoded genes contribute to evolution of heat and cold tolerance in yeast. Sci Adv 5, eaav1848 (2019).

94. Schoebel, C. N., Botella, L., Lygis, V. & Rigling, D. Population genetic analysis of a parasitic mycovirus to infer the invasion history of its fungal host. Mol Ecol 26, 2482–2497 (2017).

