## Supplementary tables and figures for "A single-stranded DNA virus replicates in the mitochondria of a marine oomycete": Supplementary files.pdf

**Table S1.** Overview of the *Halophytophthora* isolates analyzed in this study and included in the RNA pools for total RNA sequencing.

| Species | Isolate code | Pool | Site | Reference |
|---|---|---|---|---|
| *H. thermoambigua* | BD651 = CBS 147229 | Yes | Parque Natural da Ria Formosa, Santa Luzia, Tavira | Maia et al. 2022 |
| *H. thermoambigua* | BD630 | No | Rio Séqua, Tavira | Maia et al. 2022 |
| *H. thermoambigua* | BD631 | No | Rio Séqua, Tavira | Maia et al. 2022 |
| *H. thermoambigua* | BD668 | No | Ria de Alvor, Alvor, Portimão | Maia et al. 2022 |
| *H. lusitanica* | BD686 = CBS 147231 | Yes | Parque Natural da Ria Formosa, Almancil, Loulé | Maia et al. 2022 |
| *H. lateralis* | BD657 = CBS 147233 | Yes | Parque Natural da Ria Formosa, Quelfes, Olhão | Maia et al. 2022 |
| *H. frigida* | BD655 = CBS 147235 | Yes | Parque Natural da Ria Formosa, Santa Luzia, Tavira | Maia et al. 2022 |
| *H. sinuata* | BD656 = CBS 147237 | Yes | Parque Natural da Ria Formosa, Santa Luzia, Tavira | Maia et al. 2022 |
| *H. macrosporangia* | BD639 = CBS 147290 | Yes | Parque Natural da Ria Formosa, Santa Luzia, Tavira | Maia et al. 2022 |
| *H. brevisporangia* | BD662 = CBS 147238 | Yes | Parque Natural da Ria Formosa, Quelfes, Olhão | Maia et al. 2022 |
| *H. celeris* | BD885 = CBS 147240 | Yes | Parque Natural da Ria Formosa, Santa Luzia, Tavira | Maia et al. 2022 |

**Table S2.** Primers used for PCR and Sanger sequencing

| Primer name | Sequence (5’-3’) | $T_m$ |
|---|---|---|
| Halo_genomo1632V2/F | GACGACTATCCATATCTACTTGAATTTG | 58.2 |
| Halo_genomo256V2/R | AACGAGAGCTAACACTGCCATAG | 60.4 |
| Halo_genomo593V2/F | ATCTGAAATAGCTATATGTTCTTCGGATG | 59.5 |
| Halo_DNA_121 R | ACAGCAAATCCTTCAAATATACGTA | 56.9 |
| Halo_DNA_280 F | TCATTTCTGTTTTACCCATTCCAGA | 58.7 |
| Halo_DNA_536 R | AGGAGGACTAACAGCAGTAAAAGAT | 59.5 |
| Halo_DNA_780 F | CCATCCTTATGTTCTTCTTCTGCAA | 59.4 |

**Table S3.** Summary of homologous sequences identified in the mitochondrial genomes of various oomycete species, based on GenBank data.

| Organism/isolate | mtDNA accession number | mtDNA size (pb) | REP | ORF nt interval | Gemini_AL1 domain | Acc. Number | HP | ORF nt interval | domain | Acc. Number |
|---|---|---|---|---|---|---|---|---|---|---|
| *Phytophthora capsici* isolate TaiwanPc377 | BK012088 | 38,418 | N | N | N | N | Y | 6,514 - 7,627 | SMC_N super family | not annotated |
| *Phytophthora capsici* isolate TaiwanPc33e | NC_063804 | 38,427 | N | N | N | N | Y | 6,514 - 7,627 | SMC_N super family | not annotated |
| *Pseudoperonospora humuli* | MH923236 | 39,087 | N | N | N | N | orf168 | 8,382 - 7,303 | N | QLD32048.1 |
| *Pseudoperonospora cubensis* | NC_027859 | 38,995 | N | N | N | N | orf306 gene | 8,779 - 7,859 | N | YP_009164875.1 |
| *Phytophthora umbellatus* (*Phytophthora* sp. ML-2023a) | OQ735416 | 39,675 | N | N | N | N | Y | 19,330 - 18,332 | N | not annotated |
| *Paralagenidium karlingii* | NC080886 | 46,638 | N | N | N | N | ORF22 | 27,753 - 29,075 | N | YP_010886810 |
| *Paralagenidium karlingii* | NC080886 | 46,638 | N | N | N | N | ORF16 | 21,979 - 23,055 | Zinc finger-containing protein; pfam10146 | YP_010886807 |
| *Peronospora tabacina* | NC_028331 | 43,225 | orf374 | 9,988 - 11,208 | Walker A and B motifs | YP_009178776 | N | N | N | N |
| *Phytophthora botryosa* voucher CBS581_69_ORF 35_NC_067065 | NC_067065 | 39,743 | Y, | 18,882 - 17,707 | Y | not annotated | N | N | N | N |
| *Phytophthora colocasiae* voucher_Fj20_1 | OK574296 | 41,838 | Y | 19,604 - 18,456 | Y | not annotated | Y, shorter | 20,649 - 19,918 | N | not annotated |
| *Phytophthora colocasiae* isolate BC13 | BK012090 | 41,367 | Y | 8,451 - 9,599 | Y | not annotated | N | N | N | N |
| *Phytophthora colocasiae* isolate SB9 | NC_063795 | 41,297 | Y | 8,381 - 9,529 | Y | not annotated | N | N | N | N |
| *Phytophthora palmivora* | NC_056125 | 40,799 | Y | 8,338 - 9,549 | Y | not annotated | Y | 7,200 - 8,363 | SMC_prok_B super family | not annotated |
| *Lagenidium giganteum* strain ARSEF 373 | DAKRPA010000311 * | 80,403* | Y | 45,029 - 45,667 | Y | DAZ93511 | Y | 46,469 - 47,064 | Coiled-coil domain-containing protein 158; CCDC158 | DAZ93512 |
| *Phytophthora sojae* | NC_009385 | 42,977 | orf98 | 29,048 - 29,344 | Y | YP_001165402 | N | N | N | N |
| *Phytophthora sojae* | NC_009385 | 42,977 | orf112a | 29,328 - 29,666 | N | YP_001165403 | N | N | N | N |

Y, Yes, Present; N, No, absent; **Lagenidium giganteum* strain ARSEF 373 scaffold_527 mitochondrial, whole genome shotgun sequence

**Table S4**. Pairwise amino acid identity percentages among homologous replicases from distinct organisms.

| | *Phytophthora sojae* hypothetical protein_YP_001165403 | *Phytophthora palmivora* ORF_056125 | *Phytophthora colocasiae voucher Fj20_1_ORF 31_OK574296* | *Phytophthora colocasiae* isolate SB9_ORF 10_NC_063795 | *Phytophthora colocasiae* isolate BC13_ORF 21_BK012090 | *Lagenidium_giganteum*_hypothetical_protein_DAZ93511 | *Phytophthora botryosa* voucher CBS581_69_ORF 35_NC_067065 | HtCRESSV1 | *Phytophthora sojae* hypothetical protein_YP_001165402 | *Peronospora tabacina* orf374_YP_009178776 |
|---|---|---|---|---|---|---|---|---|---|---|
| *Phytophthora sojae* hypothetical protein_YP_001165403 | 0 | | | | | | | | | |
| *Phytophthora palmivora* ORF_056125 | 43.94 | | | | | | | | | |
| *Phytophthora colocasiae* voucher Fj20_1_ORF 31_OK574296 | 41.62 | 81.77 | | | | | | | | |
| *Phytophthora colocasiae* isolate SB9_ORF 10_NC_063795 | 41.62 | 82.03 | 99.74 | | | | | | | |
| Phytophthora *colocasiae* isolate BC13_ORF 21_BK012090 | 41.62 | 82.03 | 99.74 | 100.00 | | | | | | |
| *Lagenidium_giganteum*_hypothetical_protein_DAZ93511 | 42.05 | 83.26 | 98.60 | 99.07 | 99.07 | | | | | |
| *Phytophthora botryosa* voucher CBS581_69_ORF 35_NC_067065 | 41.41 | 80.40 | 79.95 | 79.69 | 79.69 | 80.00 | | | | |
| HtCRESSV1_Replicase | 38.38 | 61.19 | 61.60 | 61.60 | 61.60 | 66.98 | 60.61 | | | |
| *Phytophthora sojae* hypothetical protein_YP_001165402 | 2.13 | 30.40 | 33.49 | 33.49 | 33.49 | 2.13 | 31.02 | 27.32 | | |
| *Peronospora tabacina* orf374_YP_009178776 | 11.94 | 26.81 | 27.89 | 27.89 | 27.89 | 24.34 | 26.62 | 27.62 | 18.81 | |

| LEGEND |
|---|
| >90 % |
| 80-100 % |
| 60-80 % |
| 60-70 % |
| <60 % |

**Table S5.** Pairwise amino acid identity percentages among homologous hypothetical proteins from distinct organisms.

| | HtCRESSV1_putative_capside | *Phytophthora capsici isolate* TaiwanPc377_ORF1_BK012088 | *Phytophthora capsici* isolate TaiwanPc33e ORF1_NC_063804 | *Pseudoperonospora_humuli*_hypotehtical_protein_QLD32048.1 | *Pseudoperonospora_cubensis*_hypothetical_protein_YP_009164875.1 | *Phytophthora umbellatus*_ ORF 25_OQ735416 | *Phytophthora_colocasiae*_voucher_Fj20_1_OK574296 | *Phytophthora_palmivora*_hypothetical_protein_NC_056125 | *Lagenidium_giganteum*_hypothetical_protein_DAZ93512 | *Paralagenidium_karlingii*_YP_010886810 |
|---|---|---|---|---|---|---|---|---|---|---|
| HtCRESSV1_putative_capside | | | | | | | | | | |
| *Phytophthora capsici isolate* TaiwanPc377_ORF1_BK012088 | 58.967 | | | | | | | | | |
| *Phytophthora capsici* isolate TaiwanPc33e ORF1_NC_063804 | 58.967 | 100 | | | | | | | | |
| *Pseudoperonospora_humuli*_hypotehtical_protein_QLD32048.1 | 53.659 | 55.313 | 55.313 | | | | | | | |
| *Pseudoperonospora_cubensis*_hypothetical_protein_YP_009164875.1 | 56.875 | 58.958 | 58.958 | 69.707 | | | | | | |
| *Phytophthora umbellatus*_ ORF 25_OQ735416 | 58.808 | 59.831 | 59.831 | 60.452 | 59.935 | | | | | |
| *Phytophthora_colocasiae*_voucher_Fj20_1_OK574296 | 40.341 | 43.103 | 43.103 | 42.241 | 43.218 | 49.394 | | | | |
| *Phytophthora_palmivora*_hypothetical_protein_NC_056125 | 57.48 | 58.594 | 58.594 | 63.021 | 66.049 | 63.003 | 55.65 | | | |
| *Lagenidium_giganteum*_hypothetical_protein_DAZ93512 | 56.289 | 60.984 | 60.984 | 65.246 | 68.301 | 65.902 | 57.46 | 83.54 | | |
| *Paralagenidium_karlingii*_YP_010886810 | 21.705 | 17.373 | 17.373 | 20.043 | 21.407 | 21.409 | 17.045 | 16.564 | 20.308 | |
| *Paralagenidium_karlingii*_YP_010886807 | 23.136 | 24.084 | 24.084 | 26.053 | 25.077 | 24.865 | 18.715 | 23.951 | 26.168 | 14.768 |

**Table S6.** Confidence scores for the structural models of HP generated using AlphaFold3.

| | Ectodomain | | Full-length | |
|---|---|---|---|---|
| **Oligomeric state** | ipTM | pTM | ipTM | pTM |
| **Monomer** | - | 0.4 | - | 0.28 |
| **Dimer** | 0.08 | 0.26 | 0.22 | 0.29 |
| **Trimer** | 0.39 | 0.42 | 0.29 | 0.33 |
| **Tetramer** | 0.58 | 0.62 | 0.33 | 0.35 |
| **Pentamer** | 0.68 | 0.69 | 0.35 | 0.37 |
| **Hexamer** | 0.72 | 0.73 | 0.38 | 0.39 |
| **Heptamer** | 0.77 | 0.77 | 0.37 | 0.38 |
| **Octamer** | 0.76 | 0.76 | 0.37 | 0.38 |
| **Nonamer** | 0.74 | 0.75 | 0.35 | 0.36 |
| **Decamer** | 0.68 | 0.69 | 0.32 | 0.34 |

**Amplicon sequences from Figure S1**

>HthCRESSV1_Initial_Contig NODE_2192_length_979

CTTTATTTGTATAATTATTTTTTAATAAATTTGTTACATTATTTGTTGTAAAAATACGTAAAGTATTTTGCATAATTCGAATTGAATCTTTTATTATACGTATATTTGAAGGATTTGC
TGTATCTGAAATAGCTATATGTTCTTCGGATGATAATTCTTTTTCAAAAAATATATCATCAAAAATTACAGCTTTATGTTTTTGAGGATCATATTCACGTAATCCATGAATATTTCT
TATTAATAAAAATTTTATTCCTTGAGATTGTAAAAAGGATTTAATCATTTCTGTTTTACCCATTCCAGATAATCCATTTATTATTAATGTTTCTACACATCCTGTTGAATACCATTCT
TTTAAATCCTCTGGAATATTAAAATTTTTTACAGGTATAATAGAATTATTAAAATGATTTTCAATTTCTTTTTCATATCTAATTTTATCTATTTGTTCTAAATTTTTTAATACATTTGG
ACCTCCTTTTACCATTAACATTGGATTTGCTATTAAATGATCTTTTACTGCTGTTAGTCCTCCTTTTGCATGTAATTTAACTAAATATTCTTTAAAATCTAAATATAATTCTCCATCTA
AATATGGTAATTCCGGTTTTGCTTCAAATTTTCCTGTTTTTACAATATAACGTATTGCTGAATCTTTTTTTCGAACCGTTTCATAACTACCATGATATATTTGACCTTCTAATTCTAA
ATCTAAACAATTTGATCCATATAATTCAACTCGTTGATTAAATTCTAGAAAAACATGATAATGTTTTCCACCATCCTTATGTTCTTCTTCTGCAATTACATATTGTATTATTTTTGGT
ACTTTTTGTTCTAATTGTTCTAATATTTTTTGATTTACATTTTCTTGATTTTTTAGATTAATTTGTGAATAGGTTAAAAATAACTTTTTTGCTCTTAATCTAAATTGTTTATTTACTTTA
TTTTCTTTTTTCATATTTTAGTTTTTTAATTA

>Halo_genomo_Rv121.ab1_(reversed)

AACCATATAAAAAACCTAAAAACCAACTTTTTAACTAAAAACCAATTCTTTCTTGTGGTTTTTTATAATCTAAACCATATAAAAAACCTAAAAACCAACTTTTTAACTAAAAACCAA
TTCTTTCTTGTGGTTTTTTATAATCTAAATTTTTGAAAAATAAGATAAATACTATAAATAGCTTTTATCTTATTTTTCAATTTTATTTAAAACCAAATTAAAACGAGAGCTATGGCA
GTGTTAGCTCTCGTTTTAATAAATAAATACGCGTTAGTAATATATATATTAATTAATTTTAGATAAAAAATTATTATTATTTTGTTCTAATTCTTTTAATACTTTCATATTATGTTTAT
ATACATATTCTTCACGTTTTTTATTACGTATTTTTATTCGAACATCTATTTCAAAATCTTCTTTAAATAATGTTTTATTTATATTTAAAATAAAATATCTACTTAATACTGCTTTATTTG
TATAATTATTTTTTAATAAATTTGTTACATTATTTGTTGTAAAAATACGTAAAGTATTTTGCATAAT

>Halo_genomo_Fw780.ab1

TCCTGTTGAATACCATTCTTTTAAATCCTCTGGAATATTAAAATTTTTTACAGGTATAATAGAATTATTAAAATGATTTTCAATTTCTTTTTCATATCTAATTTTATCTATTTGTTCTA
AATTTTTTAATACATTTGGACCTCCTTTTACCATTAACATTGGATTTGCTATTAAATGATCTTTTACTGCTGTTAGTCCTCCTTTTGCATGTAATTTAACTAAATATTCTTTAAAATCT
AAATATAATTCTCCATCTAAATATGGTAATTCCGGTTTTGCTTCAAATTTTCCTGTTTTTACAATATAACGTATTGCTGAATCTTTTTTTCGAACCGTTTCATAACTACCATGATATA
TTTGACCTTCTAATTCTAAATCTAAACAATTTGATCCATATAATTCAACTCGTTGATTAAATTCTAGAAAAACATGATAATGTTTTCCACCATCCTTATGTTCTTCTTCTGCAATTAC
ATATTGTATTATTTTTGGTACTTTTTGTTCTAATTGTTCTAATATTTTTTGATTTACATTTTCTTGATTTTTTAGATTAATTTGTGAATAGGTTAAAAATAACTTTTTTGCTCTTAATCT
AAATTGTTTATTTACTTTATTTTCTTTTTTCATATTTTAGTTTTTTAATTAAAAAATAATAATTATTCATCTCCATCAAATGCATAATTATTATCTATTTTTATTTTTTTTGAAGGATTT
TCATCACTTTTCATTTCTTCGAAATCAAAAGAATCTCTTCTTCTTTTTAATATCGCTTTTTCTTCTTTTAAATTTTCAATAGATTCTGAAAAATCTACTTTATTCCAAAAATGACGACT
ATCCATATCTACTTGAATTTGATGTATTTCTTTATTTAAATTATGAATTTTTTGATCATAATTCTCTAAACGTTTATCTCGTTGTTTAATTATATTTTG

>Halo_genomo_Fw280.ab1

ACTTTTTGTTCTAATTGTTCTAATATTTTTTGATTTACATTTTCTTGATTTTTTAGATTAATTTGTGAATAGGTTAAAAATAACTTTTTTGCTCTTAATCTAAATTGTTTATTTACTTTA
TTTTCTTTTTTCATATTTTAGTTTTTTAATTAAAAAATAATAATTATTCATCTCCATCAAATGCATAATTATTATCTATTTTTATTTTTTTTGAAGGATTTTCATCACTTTTCATTTCTTC
GAAATCAAAAGAATCTCTTCTTCTTTTTAATATCGCTTTTTCTTCTTTTAAATTTTCAATAGATTCTGAAAAATCTACTTTATTCCAAAAATGACGACTATCCATATCTACTTGAATT
TGATGTATTTCTTTATTTAAATTATGAATTTTTTGATCATAATTCTCTAAACGTTTATCTCGTTGTTTAATTATATTTTGTTCTAATCTTATCTTATCTTGTTCTCTAATATATTCATCT
TTTTCTTTAATATGTTCAAATTTTTCTTTAATATGTTCAAATTTTTCAGTTTCAATCTTTTTGTTAAATTCAAACTGTTTTTGTTGAATTTGAACACTTGAATGTTGTAAATAACCTGT
GAATAAACTCCCAGCAGTTGCACTTGTTATAACTGTAATACCGTGTTTTTGCATCAAGTTTTTTACTTCATTAAAACCTATATATCGTTTTTGAATATAAAATAAAGTTTGATTTTT
ATTAAATCTTAAATTATTTATTTTTAAAATATTTTTAATTACATTATAATCTATAGTTTTACTATTATTTAAATTTTGAGGAATAAGAATGAATTTTTTGTCTATCAAATTAAATATAT
TTACTATTATTACACAGAAAAATACTAATATTGATAAACTATACGAATTATTCATTATGAGGAAGATAATGGTTAAACT

>Halo_CRESS_1632V2.ab1

ATGATTTTTTGATCATAATTCTCTAAACGTTTATCTCGTTGTTTAATTATATTTTGTTCTAATCTTATCTTATCTTGTTCTCTAATATATTCATCTTTTTCTTTAATATGTTCAAATTTTT
CTTTAATATGTTCAAATTTTTCAGTTTCAATCTTTTTGTTAAATTCAAACTGTTTTTGTTGAATTTGAACACTTGAATGTTGTAAATAACCTGTGAATAAACTCCCAGCAGTTGCAC

TTGTTATAACTGTAATACCGTGTTTTTGCATCAAGTTTTTTACTTCATTAAAACCTATATATCGTTTTTGAATATAAAATAAAGTTTGATTTTTATTAAATCTTAAATTATTTATTTTT
AAAATATTTTTAATTACATTATAATCTATAGTTTTACTATTATTTAAATTTTGAGGAATAAAAATGAATTTTTTTTCTATCAAATTAAATATATTTACTATTAATAAACAGAAAAATA
CTAATATTGATAAACTATACGAATTATTCATTATGAGGAAGATAATGTTTAAACTCCAACTTGATAATACAAATCCTGTTAAGAATACTATTAATTTTTGGTTTTTTTCATGCTCCA
CATAATTAAAATATTCAAAAATTAGTCCATATAATCCATTTTTGAATATTAAAAAATAATATAAACAAAAATTAAATAGATATGTATTTTTTATTTTATTTGTAATAAATATCCATT
GATTTATAAATAAAATTGAAATTATAATTATAATTGGATATAAATAAATCGATGGTAGTTCAAATACCACTTGATTTATTTTTATAATGATTAGATTTAAGAAAGTTATACATAAA
ATATAATTTATGTATATTTCATTAAAATTGTTTATTTTTTGTATCATTATCCAATCCTAAAAATTCTCTTATTGTATATTTAATAAATTTTATCATAATCTTTTTATCTTTCTTTTTATAT
TCAAGAGGCTAGCCTTTTACAACATTTTTATAGATATATTAAAAAGCCCTATATTTTATTTAAAACATCTTCCAAAACCCTTTTATTAAAGGTTTTCCCTAAACCATATAAAAAACC
TAAAAACCAACTTTTTAACTAAAAACC

>Halo_CRESS_256.ab1_(reversed)

GAATAAACTCCCAGCAGTTGCACTTGTTATAACTGTAATACCGTGTTTTTGCATCAAGTTTTTTACTTCATTAAAACCTATATATCGTTTTTGAATATAAAATAAAGTTTGATTTTT
ATTAAATCTTAAATTATTTATTTTTAAAATATTTTTAATTACATTATAATCTATAGTTTTACTATTATTTAAATTTTGAGGAATAAAAATGAATTTTTTTTCTATCAAATTAAATATAT
TTACTATTAATAAACAGAAAAATACTAATATTGATAAACTATACGAATTATTCATTATGAGGAAGATAATGTTTAAACTCCAACTTGATAATACAAATCCTGTTAAGAATACTATT
AATTTTTGGTTTTTTTCATGCTCCACATAATTAAAATATTCAAAAATTAGTCCATATAATCCATTTTTGAATATTAAAAAATAATATAAACAAAAATTAAATAGATATGTATTTTTT
ATTTTATTTGTAATAAATATCCATTGATTTATAAATAAAATTGAAATTATAATTATAATTGGATATAAATAAATCGATGGTAGTTCAAATACCACTTGATTTATTTTTATAATGATT
AGATTTAAGAAAGTTATACATAAAATATAATTTATGTATATTTCATTAAAATTGTTTATTTTTTGTATCATTATCCAATCCTAAAAATTCTCTTATTGTATATTTAATAAATTTTATCA
TAATCTTTTTATCTTTCTTTTTATATTCAAGAGGCTAGCCTTTTACAACATTTTTATAGATATATTAAAAAGCCCTATATTTTATTTAAAACATCTTCCAAAACCCTTTTATTAAAGGT
TTTCCCTAAACCATATAAAAAACCTAAAAACCAACTTTTTAACTAAAAACCAATTCTTTCTTGTGGTTTTTTATAATCTAAACCATATAAAAAACCTAAAAACCAACTTTTTAACTA
AAAACCAATTCTTTCTTGTGGTTTTTTATAATCTAAACCATATAAAAAACCTAAAAACCAACTTTTTAACTAAAAACCAATTCTTTCTTGTGGTTTTTTATAATCTAAATTTTTGAAA
AATAAGATAAATACTATAAATAGCTTTTATCTTA

### Supplementary Figure S1

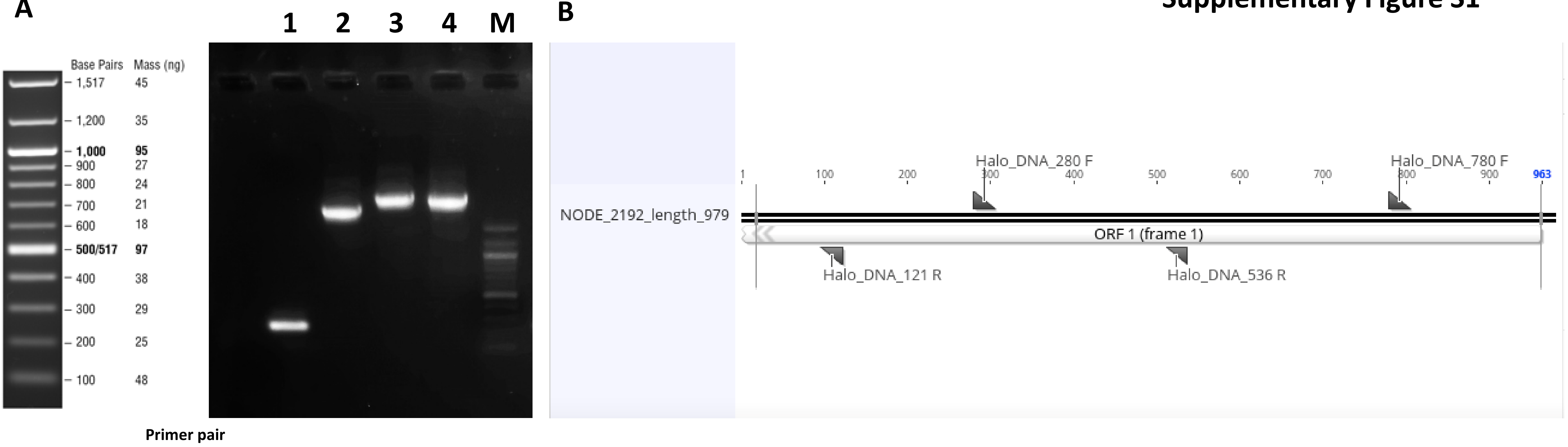


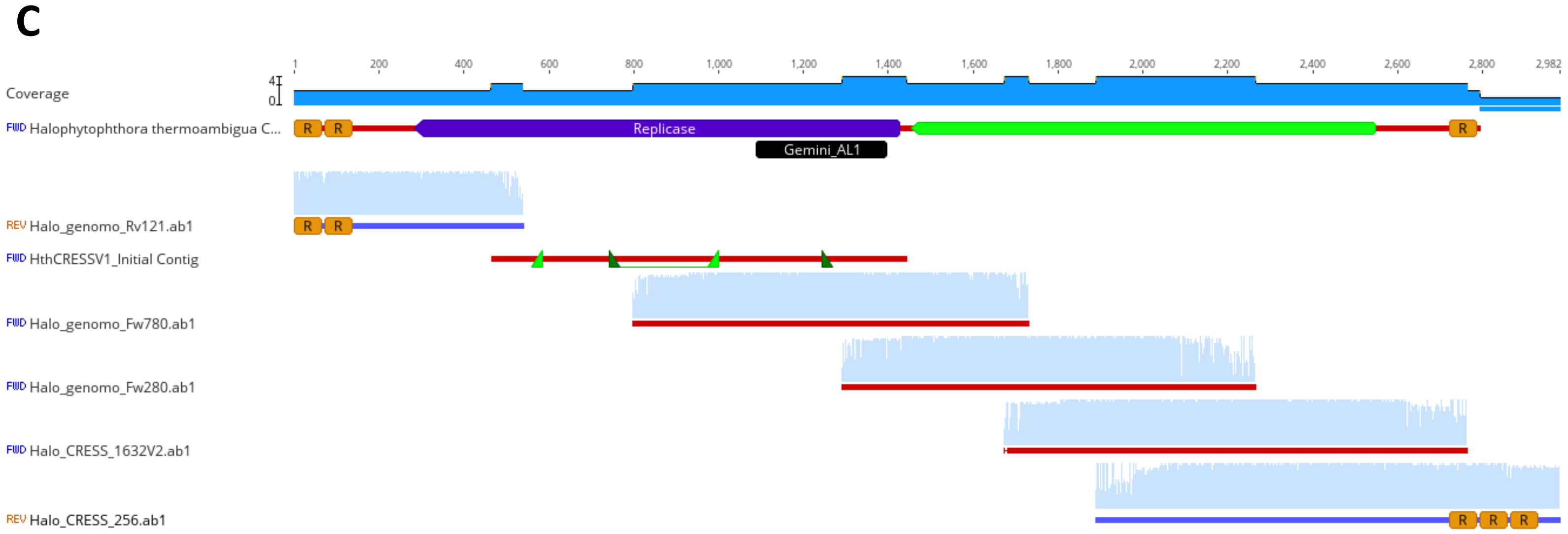


**Figure S1**. **A,** Gel picture with the amplification of the initial contig using different primer combinations: (1) Halo_DNA_280F X Halo_DNA_536R, (2) Halo_DNA_780F X Halo_DNA_121R, (3) Halo_DNA_280F X Halo_DNA_121R, (4) Halo_DNA_280F X Halo_DNA_121R (M). DNA size Marker
**B,** Picture of the primer names and sites designed in the original contig. **C,** Coverage Plots illustrating the amplicon-derived sequences in Geneious Prime. **D,** Coverage Plot illustrating the read
sense in Geneious Prime (next slide)

**Figure S1. D** Coverage Plot illustrating the read reassembling to the final sequence of HathCRESSV1 in Geneious Prime

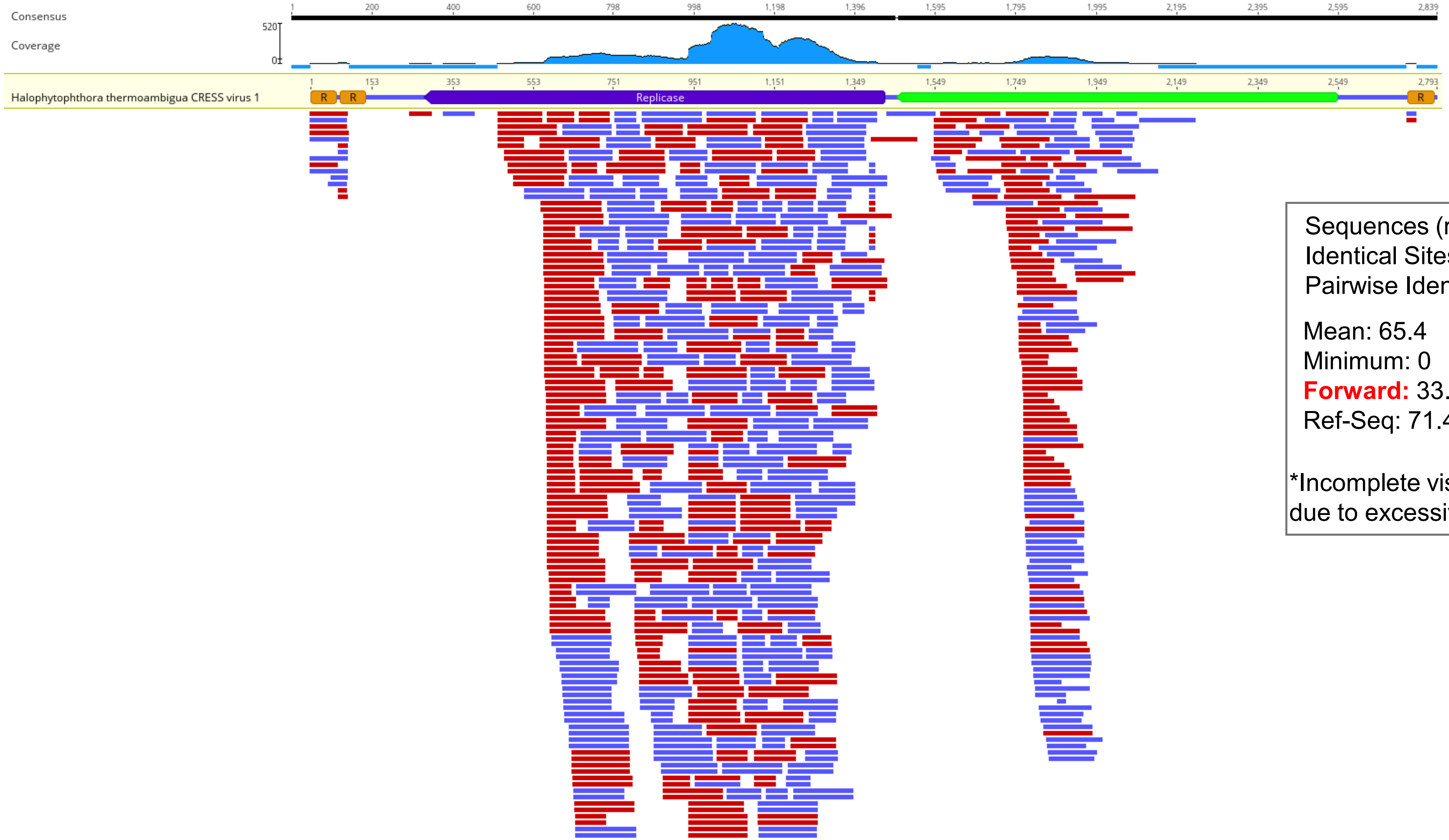


Sequences (reads): 1,637
Identical Sites: 1,847 (92.4%)
Pairwise Identity: 98.9%

| | |
|---|---|
| Mean: 65.4 | Std Dev: 116.2 |
| Minimum: 0 | Maximum: 520 |
| **Forward:** 33.3 | **Reverse:** 32.1 |
| Ref-Seq: 71.4% (1,995 of 2,793) | |

*Incomplete visualisation of the alignment due to excessive depth

**Supplementary Figure S2. Predicted secondary structures of putative stem–loop motifs in the HthCRESSV1 genome**.

**A**

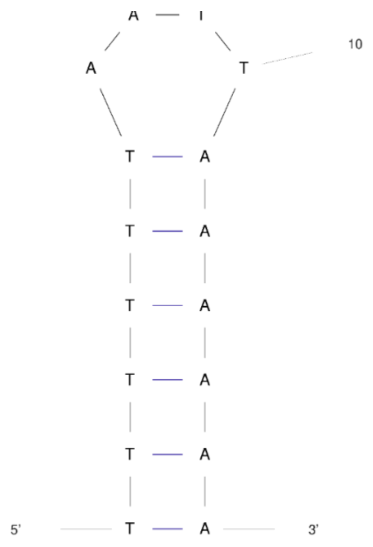


Genomic positions 1,433 – 1,448 nt. The free energy of the thermodynamic ensemble is -2.28 kcal/mol.
The frequency of the MFE structure in the ensemble is 33.15 %.
The ensemble diversity is 6.59

**B**

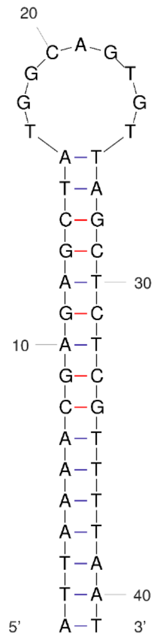


Genomic positions 214 – 254 nt. The free energy of the thermodynamic ensemble is -15.07 kcal/mol. The frequency of the MFE structure in the ensemble is 64.11 %.
The ensemble diversity is 0.86 .

**C**

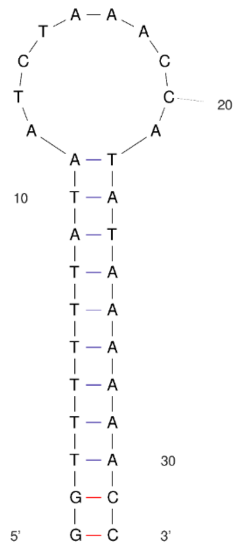


Genomic positions 57 – 88 nt and 2,778 - 2,809 nt (16 – 2,778 nt).

The free energy of the thermodynamic ensemble is -6.73 kcal/mol.
The frequency of the MFE structure in the ensemble is 49.39 %.
The ensemble diversity is 1.91 .

Secondary structures were predicted for selected genomic regions using minimum free energy folding models under DNA parameters. The analyzed regions include the SIR (**A**) and two internal genomic fragments **(B–C)**. All structures display stable stem–loop conformations, suggesting potential roles in genome replication or regulation. Nucleotide positions are indicated relative to each analyzed fragment.

**Supplementary Figure S3.** Amino acid alignment showing conserved motifs in the hypothetical protein of HthCRESSV1

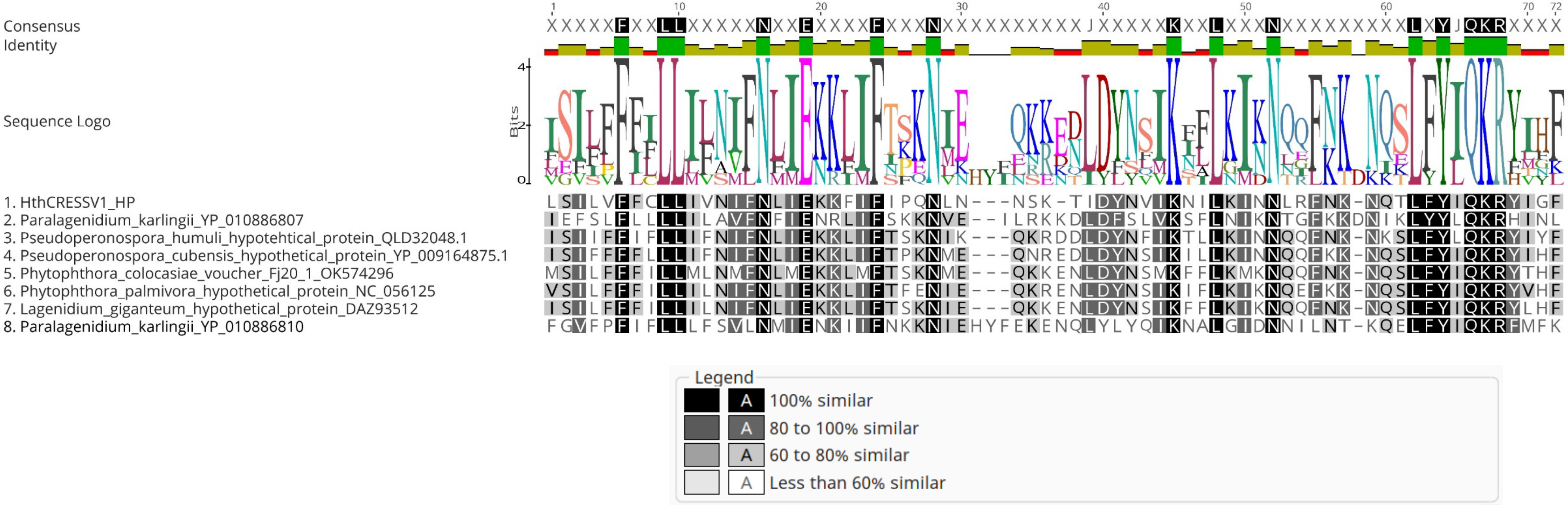

**Supplementary Figure 4S. Structural modelling of the oligomers of the HP ectodomain**

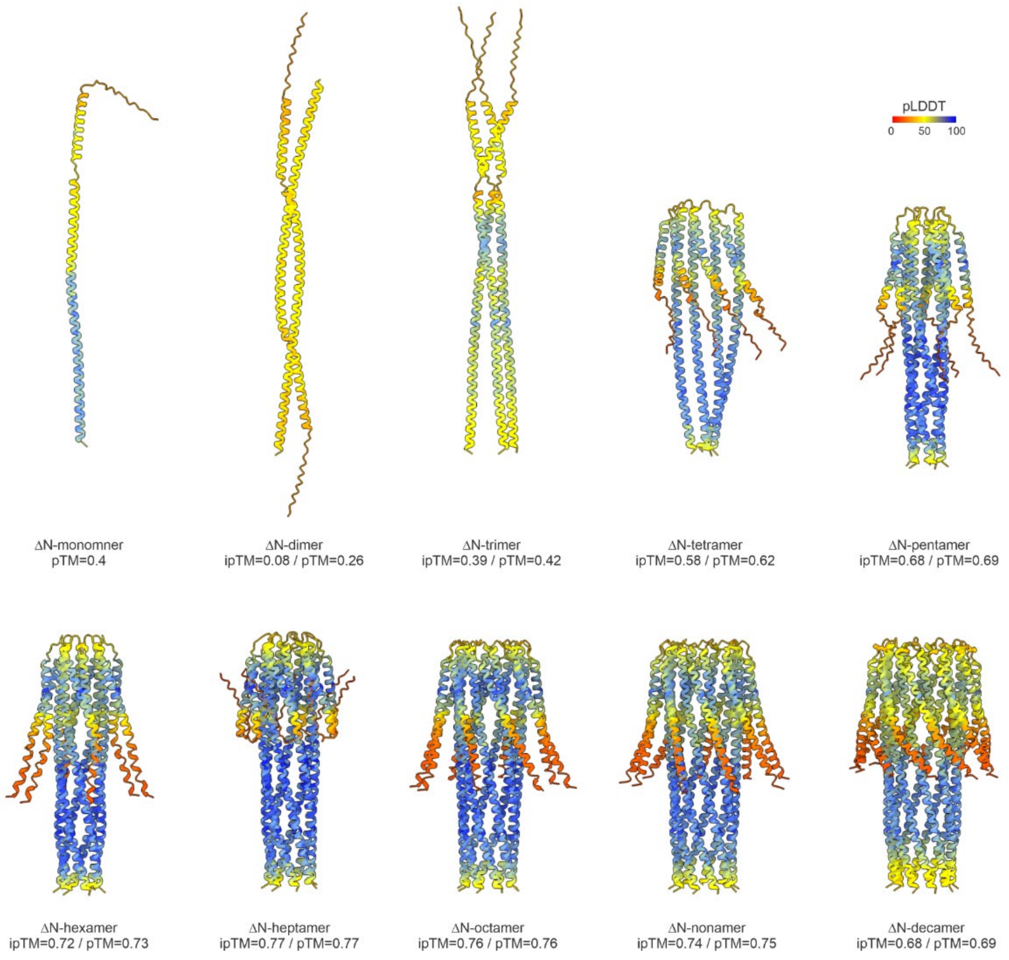


The figure shows multimeric states of the HP ectodomain from monomer to decamer colored based on the predicted local distance difference test (pLDDT) score, a per-residue measure of local confidence, from red to blue (the color scale is provided in the top right corner of the figure). The predicted template modelling (pTM) and the interface predicted template modelling (ipTM) scores are also indicated under each model.

**Supplementary Figure S5. Attempts to eliminate HthCRESSV1 from strain BD651.**

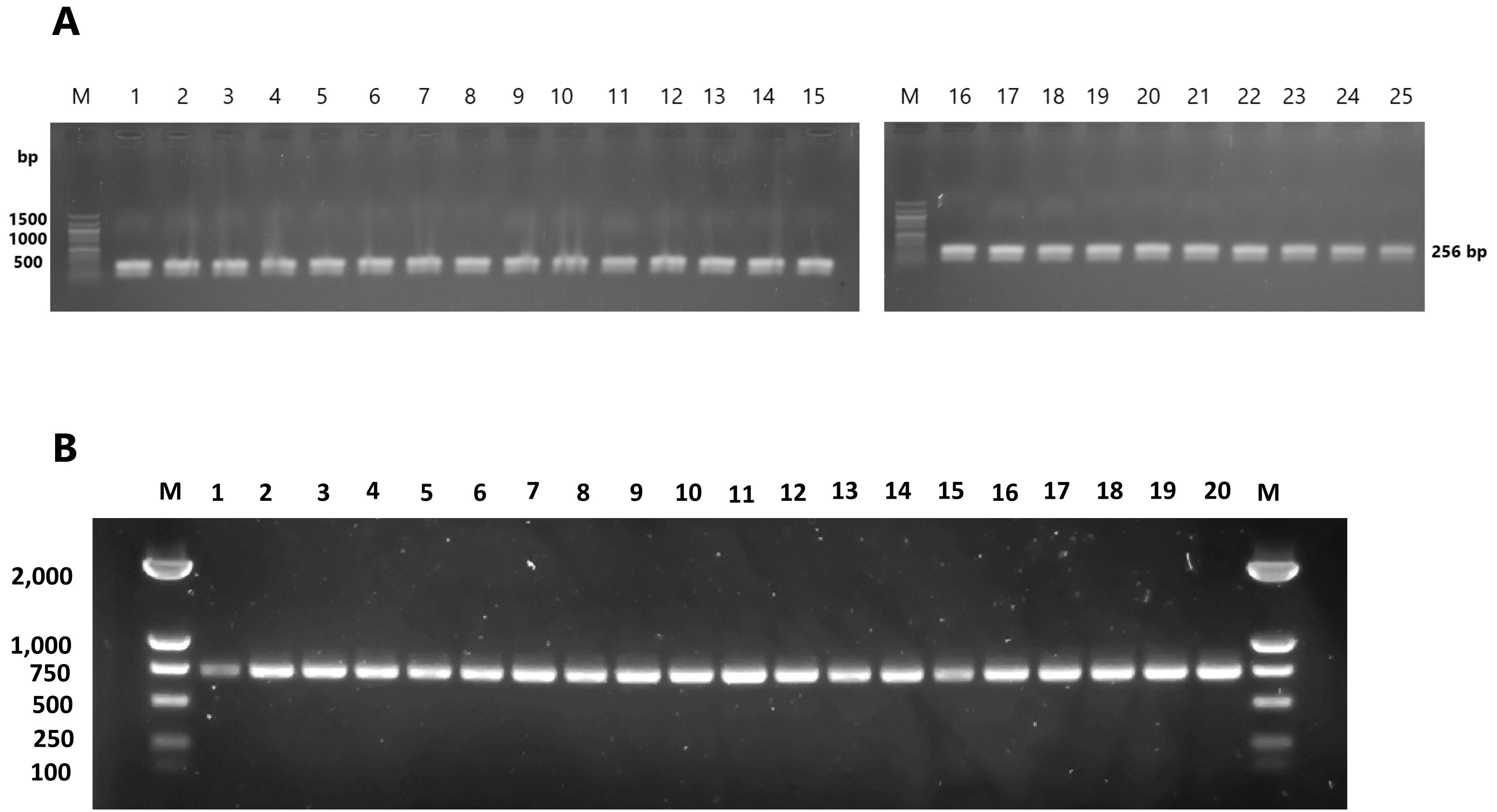


**(A)** PCR detection of HthCRESSV1 in 25 strains obtained by single zoospore isolation. M, DNA size marker. **(B)** PCR detection of HthCRESSV1 in protoplast-regenerated strains exposed to ribavirin (300 µg ml$^{-1}$ final concentration) and cycloheximide (5 µg ml$^{-1}$ final concentration) in regeneration medium. Regenerated mycelia were subsequently transferred to s. V8A medium without ribavirin and cycloheximide. M, DNA size marker.
